# Synergistic activity of the ferroptosis inducer RSL3 and Pyrimethamine to inhibit the proliferation of *Plasmodium falciparum*

**DOI:** 10.1101/2025.03.14.643256

**Authors:** Vella Nikolova, Karen Linnemannstöns, Anastasiia Zahoruiko, Markus Ganter, Carsten G. Lüder, Matthias Dobbelstein

## Abstract

Malaria tropica, caused by *Plasmodium falciparum* (*P. falciparum*), remains a global health challenge with limited therapeutic options. In mammalian cells, the small-molecule compound RSL3 induces ferroptosis via lipid peroxidation. In this study, we demonstrate that RSL3 synergizes with Pyrimethamine, an inhibitor of *P. falciparum* dihydrofolate reductase (DHFR), to suppress parasite proliferation in red blood cells (RBCs). A similar synergistic effect was observed with Cycloguanil, a structural analogue of Pyrimethamine, but not with other DHFR inhibitors or alternative ferroptosis inducers. Notably, Ferrostatin-1, an antagonist of lipid peroxidation, largely failed to rescue parasite growth in the presence of RSL3, suggesting a mechanism distinct from canonical ferroptosis. These findings suggest that the synergy may involve unidentified targets of RSL3 and Pyrimethamine in *P. falciparum*, divergent from those described in mammalian systems. Moreover, RSL3 and related compounds could serve as promising adjuvants to enhance the antimalarial efficacy of Pyrimethamine and potentially overcome drug resistance.

**Graphical abstract:** 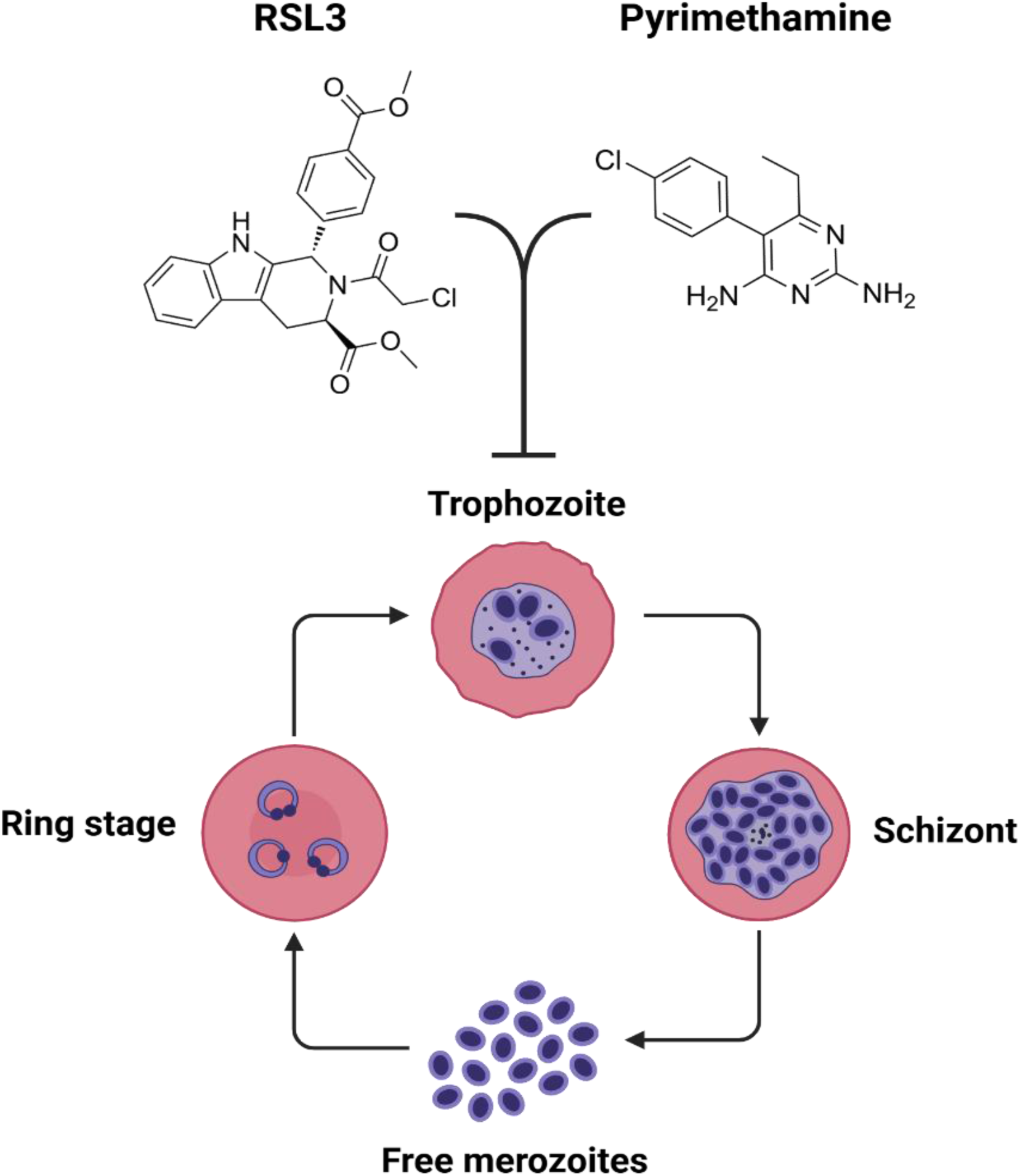

## INTRODUCTION

**Malaria** remains a significant global health challenge, claiming the lives of over half a million children under the age of five annually [1–3]. The majority of these deaths are caused by *Plasmodium falciparum (P. falciparum)*, the causative agent of malaria tropica, which is the most severe form of human malaria. While vaccination campaigns are expanding, their effectiveness is limited by the suboptimal protective efficacy of currently available vaccines. In addition to that, over the past century, numerous medications have been developed for both treatment and prevention of malaria, yet the disease persists [4]. The introduction of artemisinin and its derivatives marked a critical breakthrough in malaria therapy. However, the emergence of resistance to these drugs highlights the urgent need for novel therapeutic options [5][6]. In this context, the development of effective drug combinations is particularly valuable, as they have the potential to not only improve treatment outcomes but also slow down the emergence of resistance.

**Pyrimethamine** (marketed as Daraprim) is one of the earliest and most impactful drugs developed to combat malaria. It was synthesized in 1953 by Nobel laureates Gertrude Elion and George Hitchings at Burroughs-Wellcome (now part of GlaxoSmithKline) [7], and remains on the World Health Organization’s (WHO) list of essential medications to date. Its simple chemical structure enables economical and straightforward synthesis. Pyrimethamine continues to be used, particularly in sub-Saharan Africa, for intermittent preventive treatment during pregnancy, where it helps reduce the risk of low birth weight associated with malaria [8–10], but also for seasonal malaria chemoprevention among children [11]. A related antimalarial drug – Proguanil, which is metabolized into Cycloguanil – is primarily used for prophylaxis [12].

Both Pyrimethamine and Cycloguanil target *P. falciparum* dihydrofolate reductase (DHFR), an enzyme with a crucial role in the folate pathway [13]. DHFR is responsible for regenerating tetrahydrofolate (THF), which is essential for one-carbon (C1) transfer reactions, such as the conversion of deoxyuridine monophosphate (dUMP) to deoxythymidine monophosphate (dTMP). dTMP is subsequently phosphorylated to form deoxythymidine triphosphate (dTTP), a key building block in DNA synthesis. The conversion of dUMP to dTMP generates dihydrofolate (DHF), which is recycled back to THF by DHFR to sustain DNA synthesis. By inhibiting DHFR, Pyrimethamine and Cycloguanil disrupt DNA synthesis, thereby suppressing the replication of *P. falciparum*.

Despite its efficacy, Pyrimethamine is prone to resistance development [14, 15]. Resistance typically arises from specific mutations in *P. falciparum* DHFR, which reduce the drug’s binding affinity while retaining its enzymatic activity. This presents a significant challenge for the clinical use of Pyrimethamine. While the drug remains effective in regions where DHFR mutations have not fully compromised its activity, its vulnerability to resistance highlights the need for a’hit hard and early’ approach to minimize resistance development. Drug combinations offer a promising solution to this challenge. Currently, Pyrimethamine is often paired with sulfonamides, such as Sulfadoxine, which inhibit dihydropteroate synthase (DHPS), an enzyme involved in folate precursor synthesis. This drug combination, sold as Fansidar, has been widely used but is also hampered by resistance. Consequently, there is an ongoing need to identify and develop new drug combinations, in order to enhance the antimalarial efficacy of Pyrimethamine while mitigating resistance.

**Ferroptosis** is a distinct form of programmed cell death driven by the accumulation of toxic amounts of lipid peroxides [16, 17]. It has been primarily studied as a mechanism to eliminate cancer cells [18, 19]. This process involves a series of redox reactions, including the Fenton reaction, which generates lipid peroxides in cellular membranes and ultimately leads to cell death. Ferroptosis can be induced by a number of compounds through various mechanisms. For instance, Erastin and its analogue Erastin2 block the cellular uptake of cystine, thereby impeding the synthesis of glutathione [20], which is one of the major cellular antioxidants. Conversely, ferroptosis can be inhibited by molecules that act as radical-trapping antioxidants (RTAs), such as Ferrostatin-1 [21].

Among ferroptosis inducers, the small-molecule compound RAS-selective lethal 3 (RSL3) has been identified as particularly effective. RSL3 contains a chloroacetamide moiety, and such moieties are capable of covalently modifying thiol groups on cysteine residues or imidazole rings within histidine residues [22–25]. Its primary target was reported to be glutathione peroxidase 4 (GPX4), a phospholipid hydroperoxidase that prevents lipid peroxidation [26]. However, additional targets have been proposed to contribute to the ferroptosis-inducing activity of RSL3, including thioredoxin reductase 1 (TXNRD1) [27] and other selenoproteins [28].

**Combining RSL3 with Pyrimethamine**: Recent findings have shown that RSL3 interferes with the proliferation of *P. falciparum in vitro* [29], suggesting that RSL3 and other ferroptosis inducers could have potential as antimalarial agents. Building on this premise, we aimed to explore combinations of RSL3 with established antimalarial drugs to enhance its efficacy. Among the candidates, dihydrofolate reductase (DHFR) inhibitors, such as Pyrimethamine, emerged as promising partners. DNA synthesis in *P. falciparum* requires DHFR for production of thymidine deoxynucleotide, and also depends on reducing equivalents for ribonucleotide reduction – processes that are disrupted under oxidative stress. Supporting this rationale, prior studies have demonstrated that inhibiting glutathione biosynthesis sensitizes the rodent malaria species *Plasmodium berghei* to antifolates [30]. Furthermore, RSL3 has been shown to target thioredoxin reductase 1 (TXNRD1), an enzyme critical for ribonucleotide reduction via its role in regenerating thioredoxin [27], raising the perspective that RSL3 and Pyrimethamine might target DNA synthesis from different angles, thus synergizing in suppressing the growth of *P. falciparum*.

In this study, we demonstrate that Pyrimethamine and its relative Cycloguanil exhibit strong synergy with RSL3 in the elimination of *P. falciparum* from cultures. Initially, we hypothesized that this synergistic effect was primarily mediated by the ferroptosis-inducing activity of RSL3. However, this mechanistic concept was challenged, as other ferroptosis inducers and alternative DHFR inhibitors did not achieve a comparable degree of synergy. Despite this complexity, the combination of RSL3 with Pyrimethamine emerges as a novel and promising antimalarial strategy that warrants further exploration and development.

## RESULTS

### RSL3 strongly synergizes with Pyrimethamine to inhibit the proliferation of *P. falciparum* within human RBCs

RSL3 is an inducer of lipid peroxidation in mammalian cells. This prompted us to test whether RSL3 might synergize with traditional antimalarial drugs, such as Dihydroartemisinin and Atovaquone, which often act by conferring oxidative stress to the parasites. In addition to that, we also tested another commonly used antimalarial – Pyrimethamine, which inhibits *P. falciparum* DHFR. To assess the antimalarial activity of the drugs, we treated asynchronously growing *P. falciparum* parasites, strain 3D7, with varying concentrations of RSL3, Dihydroartemisinin, Atovaquone and Pyrimethamine, either individually or in combination. Parasite growth was quantified by SYBR Green staining of DNA. As expected, RSL3 and Dihydroartemisinin suppressed parasite growth at low micromolar concentrations, while Atovaquone and Pyrimethamine were active even in the nanomolar range (Figure 1, A-C, and Suppl. Fig. 1, 2 and 3). However, when combining RSL3 with either Dihydroartemisinin (Figure 1A, Suppl. Fig. 1) or Atovaquone (Figure 1B and Suppl. Fig. 2), no synergistic activity was observed.

**Figure 1:**
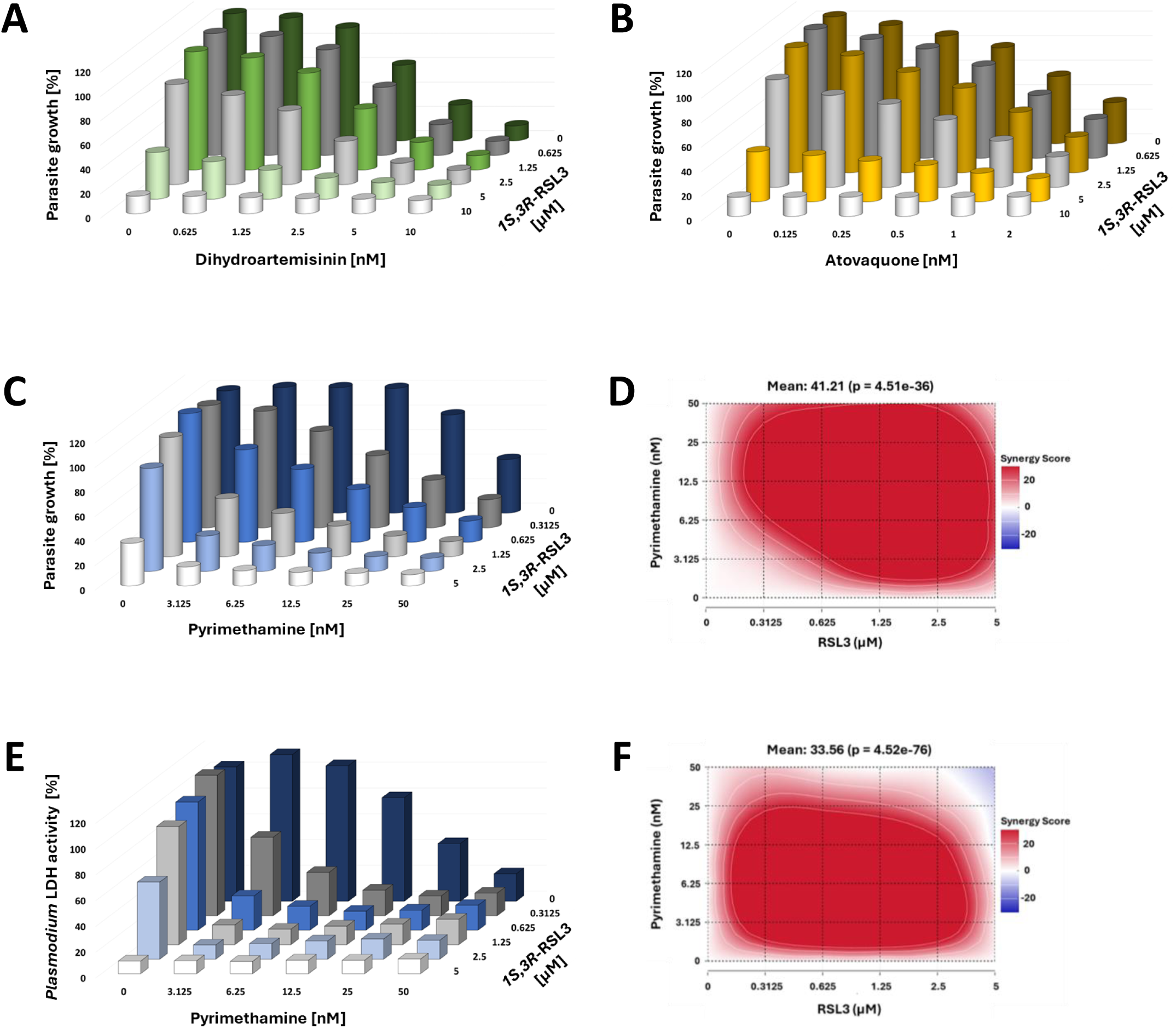
Synergy between RSL3 and Pyrimethamine against *P. falciparum*, in contrast to Dihydroartemisinin and Atovaquone. **A.** Asynchronous *P. falciparum* parasites, strain 3D7, were treated with RSL3 (isoform *1S,3R*-RSL3) and/or Dihydroartemisinin (DHA) at the indicated concentrations for 72 hours, followed by measurement of the total DNA content by SYBR Green fluorescence. Since the host RBCs lack endogenous DNA, the signal is proportional to the DNA content of the parasites within the cultures. All values were normalized to the values measured in the absence of any drugs. Suppl. Fig. 1 represents the values obtained from each biological replicate, which were plotted individually for each concentration of DHA. **B.** Analogous experiments were carried out as in (**A**), but parasites were treated with RSL3 and/or Atovaquone. Suppl. Fig. 2 shows the values from each biological replicate, which were plotted individually for each concentration of Atovaquone. **C.** Parasites were treated with RSL3 and/or Pyrimethamine, as described in (**A**). Suppl. Fig. 3 shows the values from each biological replicate, which were plotted individually for each concentration of Pyrimethamine. **D.** Bliss synergy scores were determined from the data shown in (**C**) to quantify the degree of synergy between the two drugs. **E.** Analogous experiments were carried out as in (**C**), but using *P. falciparum* LDH activity as another readout that corresponds to the growth of the parasites. Suppl. Fig. 4 shows the values from each biological replicate, which were plotted individually for each concentration of Pyrimethamine. **F.** Bliss synergy scores were calculated based on the LDH activities from (**E**), as in (**D**).

Strikingly, when RSL3 and Pyrimethamine were combined, they interfered with parasite growth synergistically. As single drugs, 27.9 nM pyrimethamine or 4.29 µM RSL3 were required to achieve 50% inhibition of *P. falciparum* growth. In contrast, when the drugs were combined, much lower concentrations were sufficient to produce a comparable effect. For instance, a combination of 6.25 nM Pyrimethamine with 1.25 µM RSL3 significantly inhibited the proliferation of *P. falciparum* (Figure 1C, Supplemental Figure 3). The synergistic interaction between RSL3 and Pyrimethamine was further quantified by calculation of the Bliss synergy scores [31], which exceeded 20 over a broad range of drug concentrations (Figure 1D). Notably, a Bliss score of 10 is already considered indicative of synergy. These findings were corroborated using an alternative readout - the measurement of *P. falciparum* lactate dehydrogenase (LDH) levels [32], which showed a similar pattern of strong synergy between the two drugs (Figure 1, E and F, Supplemental Figure 4). In summary, RSL3 and Pyrimethamine exhibit a powerful synergistic effect in inhibiting the proliferation of *P. falciparum* in human RBCs, highlighting their potential as a combinatorial therapeutic strategy.

### The combination of RSL3 and Pyrimethamine suppresses the growth of *P. falciparum* sustainably, even after drug washout

To evaluate the sustainability of the combination between RSL3 and Pyrimethamine, asynchronously growing parasites were treated with the drugs either individually or in combination. After 48 hours, the drugs were washed off and the cultures were incubated further. Parasitemia was assessed at various time points using Giemsa staining. Transient drug exposure failed to suppress *P. falciparum* growth even when high concentrations of RSL3 (5 µM) or Pyrimethamine (50 nM) were used as single treatments (Figure 2, A and B, and Suppl. Fig. 5, A, B, D, E). In contrast, the combination of RSL3 (1.25 µM) and Pyrimethamine (25 nM) eliminated detectable parasites completely, with no resurgence of *P. falciparum* growth observed two days after drug washout (Figure 2C, Suppl. Fig. 5, C and F). Within the limitations of the observation period, these results indicate that the combination of RSL3 and Pyrimethamine has a significantly more sustainable inhibitory effect on *P. falciparum* than either drug alone.

**Figure 2:**
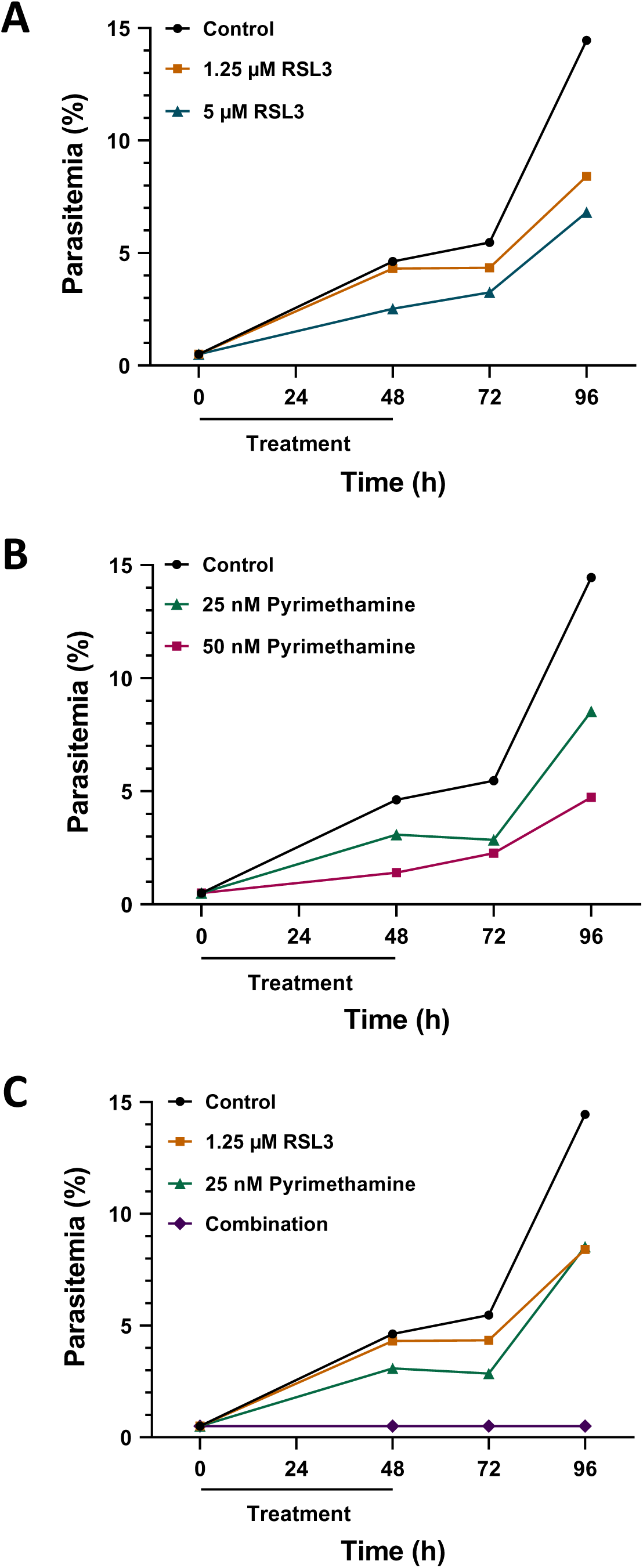
Sustainable suppression of *P. falciparum* growth by RSL3 and Pyrimethamine. **A.** *P. falciparum* cultures were synchronised using 5% D-sorbitol. Ring-stage parasites (0.5% parasitemia) were treated with the indicated concentrations of RSL3. After 48 hours of treatment, the drugs were washed off and plasmodia were incubated further. At the indicated time points, parasitemia was determined by Giemsa staining. A total of approx. 3000 parasites were counted from three biological replicates, and a representative replicate is shown here. Suppl. Fig. 5 displays the other two biological replicates. **B.** Analogous experiments as in (**A**) were carried out using Pyrimethamine. **C.** RSL3 and Pyrimethamine were combined in a setting as in (**A**) and (**B**).

**Figure 3:**
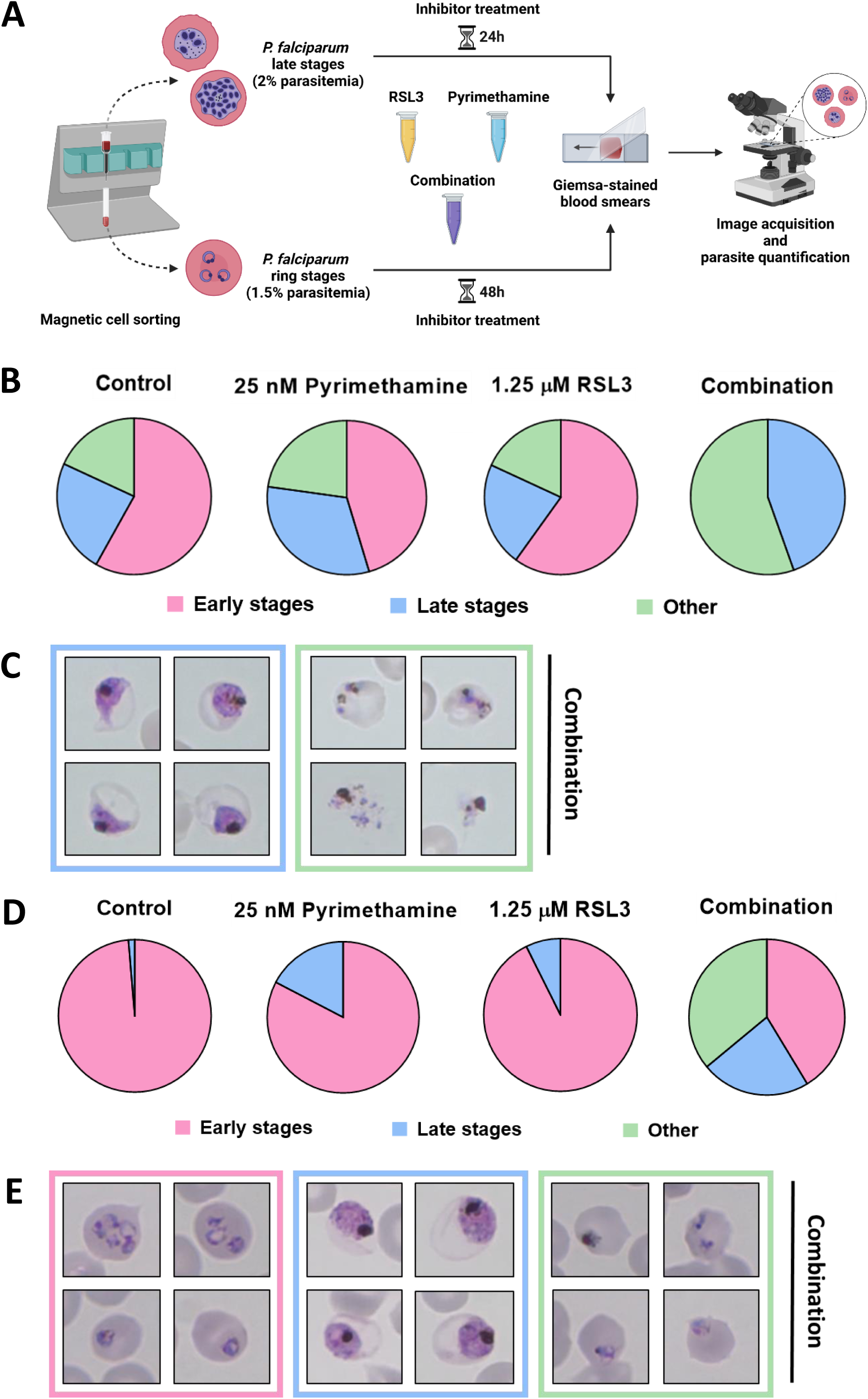
**Activity of RSL3 and Pyrimethamine on late blood stages of *P. falciparum* A.** Experimental setup for synchronisation of *P. falciparum* parasites through magnetic cell sorting (MACS). Ring stages obtained from the flow-through were treated with RSL3, Pyrimethamine, or the combination, followed by Giemsa staining after another 48 hours. Again, through MACS, late-stage parasites were enriched and eluted from the magnetic column. These were treated with RSL3 and Pyrimethamine individually or in combination, followed by Giemsa staining after another 24 hours. **B.** Quantification of the stage distribution of plasmodia after treatment of ring-stage parasites (pie charts). RBCs were classified according to the plasmodia they contained, either early or late stages, or displaying morphologies that could not be attributed to a particular stage or were not normally found in *P. falciparum* cultures (labelled as “other”). Quantification is based on counting a total of 110 parasites. **C.** Representative images of late-stage parasites (framed in blue) and parasites displaying abnormal morphologies (framed in green) after combinatorial treatment with RSL3 and Pyrimethamine, corresponding to (**B**). **D.** Stage distributions from treatment of late-stage parasites as described in (**A**), displayed as in (**B**). Quantification is based on counting a total of 150 parasites. **E.** Representative images of ring stages (framed in pink), late-stage parasites (framed in blue) and parasites displaying abnormal morphologies (framed in green) after combinatorial treatment with RSL3 and Pyrimethamine, corresponding to (**D**).

### The combination of RSL3 and Pyrimethamine prevents the subsequent round of invasion

To determine the stage at which RSL3 and Pyrimethamine interfere with the growth of *P. falciparum*, synchronized parasites were treated and analysed by Giemsa staining. Ring-stage parasites were obtained from the flow-through after magnetic sorting [33], and were incubated for 48 hours in the presence or absence of the drugs, individually or in combination (Figure 3A). In untreated cultures, most parasites progressed through their developmental cycle and returned to the ring stage. Similarly, treatment with either RSL3 (1.25 µM) or Pyrimethamine (25 nM) alone only marginally altered this progression. In contrast, the drug combination virtually eliminated ring stages and arrested parasites predominantly at the trophozoite-to-schizont transition, occasionally displaying an unusual morphology (Figure 3, B and C). These findings suggest that the combination of RSL3 and Pyrimethamine prevents the development of mature schizonts, which are essential for the subsequent invasion of new RBCs.

To further confirm this, late-stage *P. falciparum* parasites (trophozoites and schizonts) were enriched by magnetic sorting and incubated for 24 hours in the presence or absence of the drugs, both individually or in combination (Figure 3A). In untreated cultures, most parasites successfully progressed to the ring stage. Single-drug treatments still permitted the majority of parasites to complete this transition. However, the drug combination drastically reduced the number of parasites reaching the ring stage, with a higher fraction arrested at late stage instead (Figure 3, D and E). In addition to that, a considerable fraction of parasites displayed abnormal morphologies, which could not be assigned to any stage. In summary, these results strongly suggest that the combination of RSL3 and Pyrimethamine primarily targets *P. falciparum* in its late blood stages, effectively preventing subsequent RBC invasion and parasite proliferation.

### Cycloguanil exhibits antiplasmodial synergy with RSL3, comparable to Pyrimethamine

Next, we investigated whether Pyrimethamine could be replaced by similar compounds to achieve synergistic effects with RSL3. Cycloguanil, a well-known antimalarial and the active metabolite of the drug Proguanil (marketed as Paludrine), is the closest structural relative of Pyrimethamine (Figure 4A). It acts as an inhibitor of *P. falciparum* DHFR [34], although with subtle structural differences in binding [35]. Proguanil is widely used for malaria prophylaxis, particularly in combination with Atovaquone, sold under the brand name Malarone [36].

**Figure 4:**
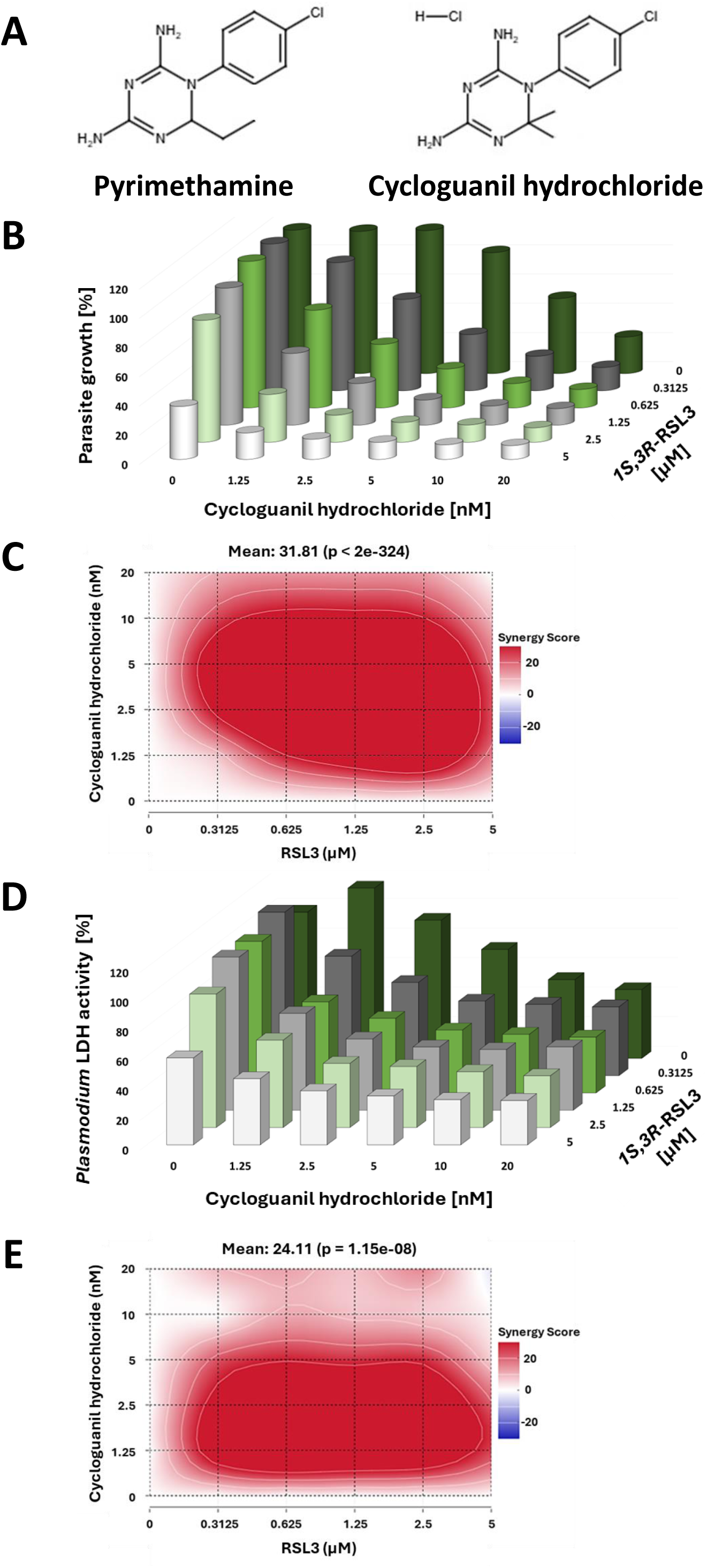
Synergy between RSL3 and Cycloguanil. **A.** Structures of Pyrimethamine and Cycloguanil hydrochloride, visualizing their similarities. Structures were generated using the *Chemical Sketch* software provided by www.rcsb.org. **B.** Asynchronously growing *P. falciparum* parasites were incubated with RSL3 and/or Cycloguanil for 48 hours, followed by DNA quantification using SYBR Green staining. All data are indicated as percentages of the value obtained in the absence of any drugs. Suppl. Fig. 6 displays the values from each biological replicate, which were plotted individually for each concentration of Cycloguanil. **C.** Bliss synergy scores corresponding to (**B**) to indicate the degree of synergy. **D.** As in (**A**) but using *P. falciparum* LDH activity as a readout. Suppl. Fig. 7 displays the values from each biological replicate, which were plotted individually for each concentration of Cycloguanil. **E.** Bliss synergy scores corresponding to the graph in (**D**).

When RSL3 was combined with Cycloguanil in the treatment of asynchronously growing *P. falciparum* parasites, we observed strong synergies comparable to those seen with Pyrimethamine. This was consistent across two independent readouts: SYBR-Green staining of parasite DNA and *P. falciparum* LDH quantification (Figure 4, B–E, Suppl. Fig. 6 and 7).

In conclusion, Cycloguanil and Pyrimethamine exhibit similar synergistic interactions with RSL3, effectively enhancing the elimination of *P. falciparum* within human RBCs.

### RSL3 does not synergize with the DHFR inhibitors WR99210 and Methotrexate

To further explore the potential of different antimalarial combinations with RSL3, we assessed the impact of additional DHFR inhibitors on *P. falciparum in vitro*. First, we evaluated WR99210 – a specific inhibitor of *P. falciparum* DHFR [37], which is commonly used for selecting recombinant *P. falciparum* strains with human DHFR as a selection marker [38]. Additionally, we tested Methotrexate – one of the oldest anticancer drugs, which acts as a competitive inhibitor of human DHFR [39] and also inhibits *P. falciparum* DHFR [40]. Surprisingly, neither of the compounds exhibited synergistic activity when combined with RSL3 (Figure 5, A and B, Suppl. Fig. 8 and 9). While each compound demonstrated antiplasmodial activity individually, their combinations with RSL3 showed no enhancement in efficacy. These findings strongly suggest that DHFR inhibition alone does not underpin the observed synergy between RSL3 and Pyrimethamine or Cycloguanil. Instead, other mechanisms or molecular targets of these diamino compounds likely contribute to their cooperative interactions with RSL3.

**Figure 5:**
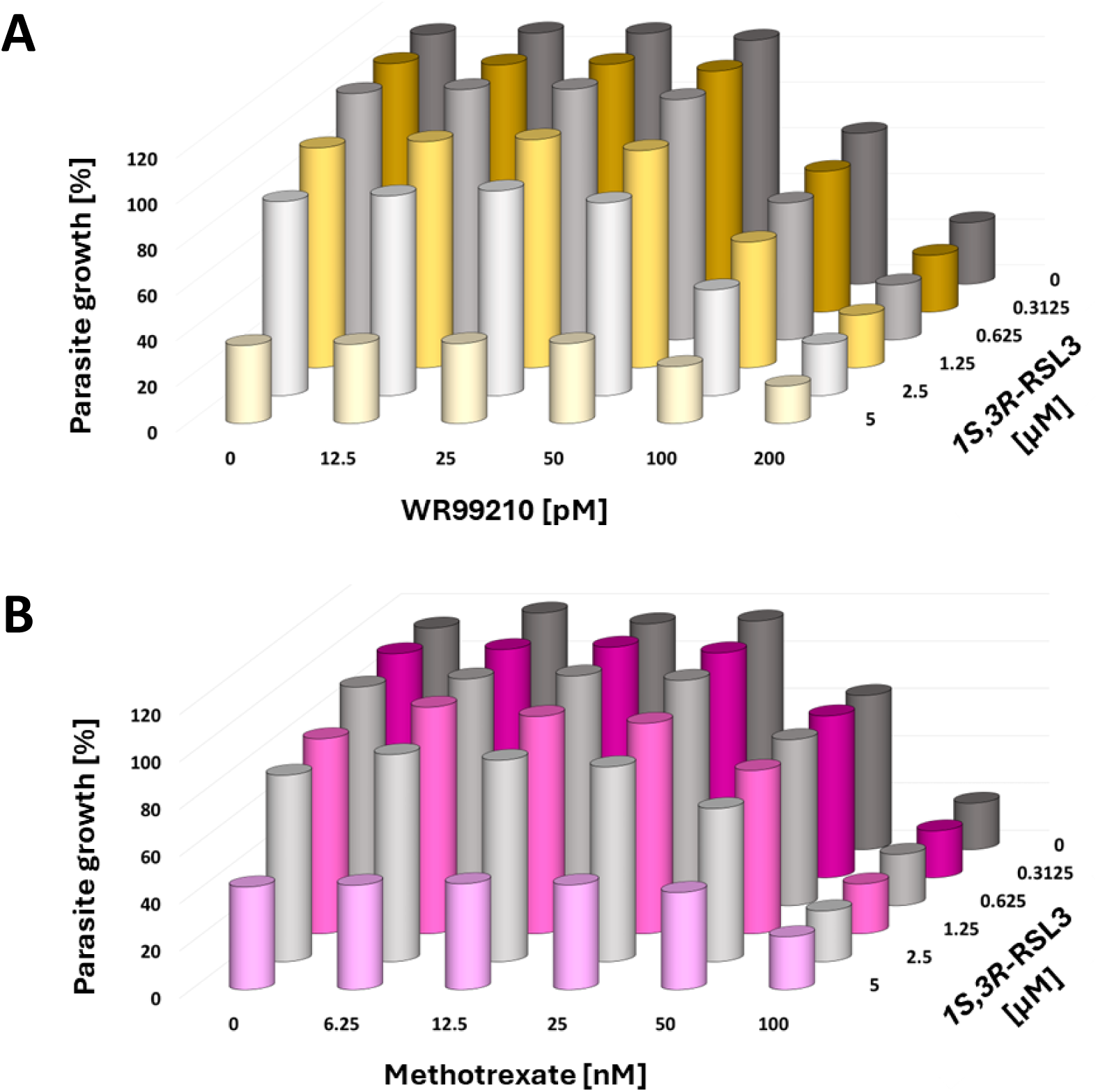
Lack of cooperation between RSL3 and WR99210 or Methotrexate. **A.** Parasites were treated with RSL3 and the *P. falciparum* DHFR inhibitor WR99210 for 48 hours, followed by quantification of DNA through SYBR Green staining. No indication of synergy was observed. Suppl. Fig. 8 displays the values from each biological replicate, which were plotted individually for each concentration of WR99210. **B.** Combination of RSL3 with the general DHFR inhibitor Methotrexate. Again, no synergistic effects were observed between these two drugs. Suppl. Fig. 9 displays the values from each biological replicate, which were plotted individually for each concentration of Methotrexate.

### The epimer *1R,3R*-RSL3 compromises *P. falciparum* proliferation more strongly than the commonly used *1S,3R*-RSL3, but shows a lower degree of synergy with Pyrimethamine

To investigate whether RSL3 could be replaced by structurally related compounds to achieve synergy with Pyrimethamine, we compared the activity of two RSL3 stereoisomers. RSL3 contains two chiral centers, resulting in stereoisomers with distinct configurations (Figure 6A). The most commonly used form is *1S,3R*-RSL3, which has been employed in all experiments described, unless stated otherwise. This isomer was previously identified as a specific ligand of the enzyme GPX4. In contrast, its epimer, *1R,3R*-RSL3, differs in the configuration at one chiral center and has been reported to be less active against GPX4 while retaining similar overall chemical properties [26].

**Figure 6:**
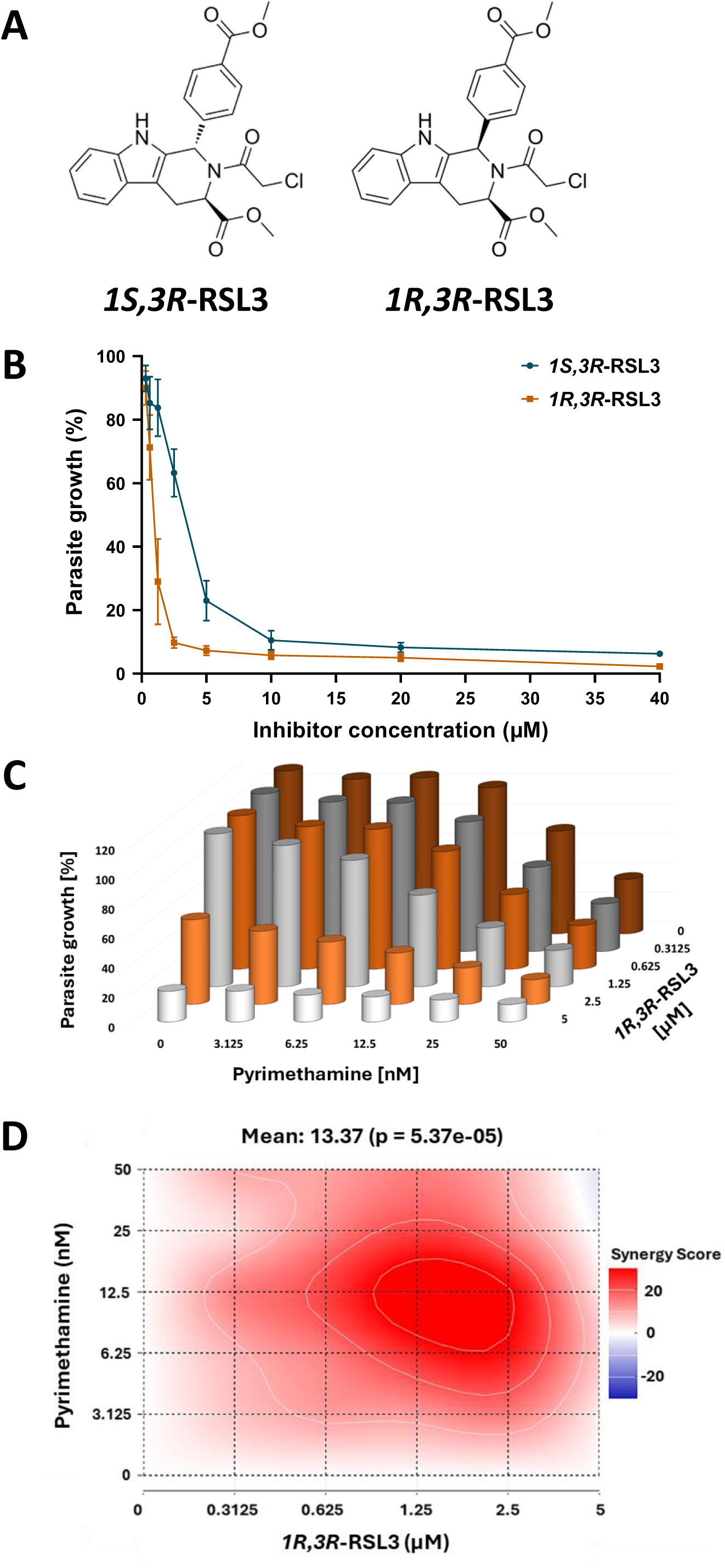
**Activity of *1R,3R*-RSL3 against *P. falciparum* A.** Structures of RSL3 stereoisomers. *1R,3R*-RSL3 is an epimer of the most widely used GPX4 inhibitor *1S,3R*-RSL3. According to its initial description [26], *1S,3R*-RSL3 has a far stronger affinity to GPX4 than *1R,3R*-RSL3. **B.** Activity of *1R,3R*-RSL3 against *P. falciparum*, in comparison to the *1S,3R*-RSL3 isomer, which was used in all other experiments shown here. Asynchronously growing *P. falciparum* parasites were treated with the two compounds at various concentrations for 72 hours, followed by assessment of the total DNA levels in comparison to non-treated cultures. Raw data on the impact of *1S,3R*-RSL3 and *1R,3R*-RSL3 on the intraerythrocytic proliferation of *P. falciparum* is shown in Suppl. Fig. 10. **C.** The isomer *1R,3R*-RSL3 was combined with Pyrimethamine as indicated, followed by determination of the total DNA content. Suppl. Fig. 11 displays the values from each biological replicate, which were plotted individually for each concentration of Pyrimethamine. **D.** Bliss synergy scores corresponding to the graph in (**C**). For comparison with *1S,3R*-RSL3, please see Figure 1, C and D, and Suppl. Fig. 3.

**Figure 7:**
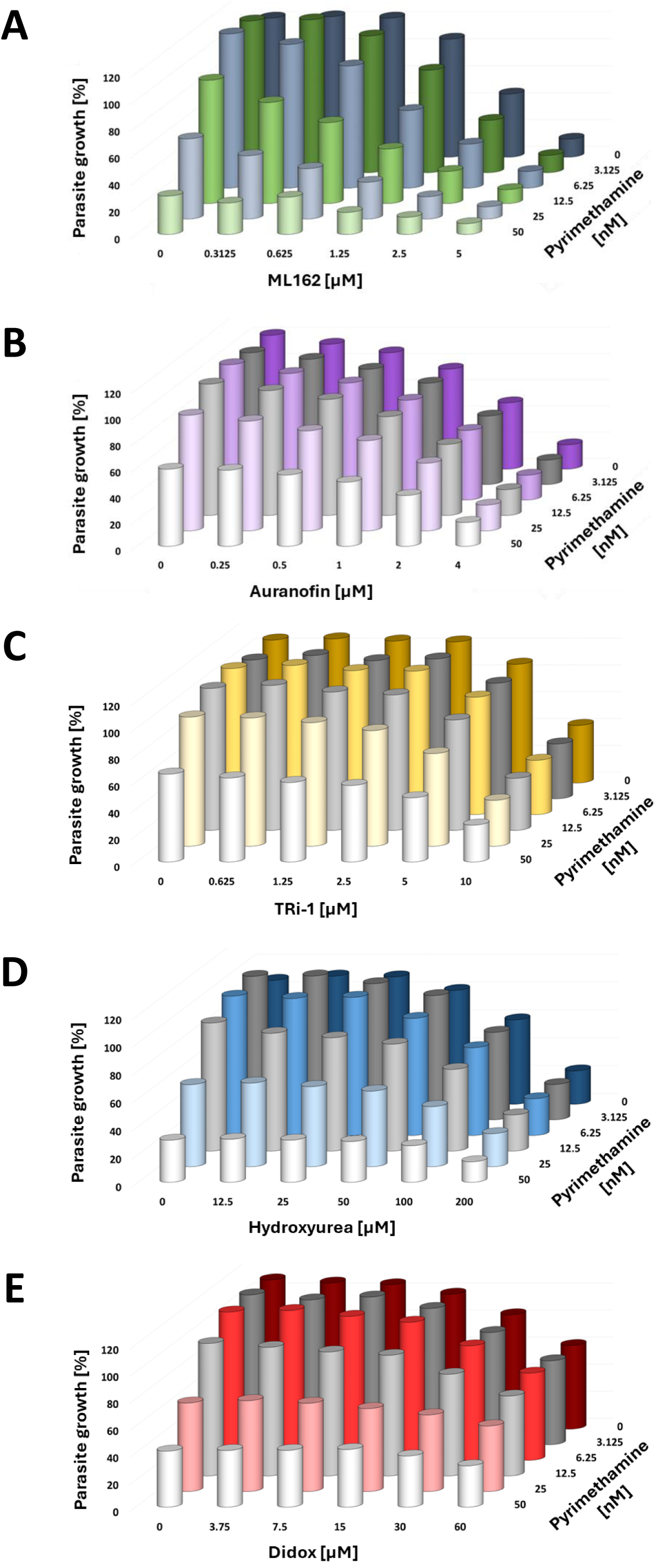
Lack of synergy between Pyrimethamine and inhibitors of thioredoxin reductase or ribonucleotide reductase. **A.** ML162, an inhibitor of thioredoxin reductase, was combined with Pyrimethamine to treat *P. falciparum*, followed by quantification of DNA to assess the relative suppression of parasite growth. Suppl. Fig. 12 displays the values obtained from each biological replicate, which were plotted individually for each concentration of ML162. **B.** Auranofin (inhibitor of thioredoxin reductase) with Pyrimethamine. Suppl. Fig. 13 displays the values obtained from each biological replicate, which were plotted individually for each concentration of Auranofin. **C.** TXNRD1 inhibitor 1 (TRi-1) with Pyrimethamine. Suppl. Fig. 14 displays the values obtained from each biological replicate, which were plotted individually for each concentration of TRi-1. **D.** Hydroxyurea (ribonucleotide reductase inhibitor) with Pyrimethamine. Suppl. Fig. 15 displays the values obtained from each biological replicate, which were plotted individually for each concentration of Hydroxyurea. **E.** Didox (3,4-dihydrosybenzohydroxamic acid; inhibitor of ribonucleotide reductase) with Pyrimethamine. Suppl. Fig. 16 displays the values obtained from each biological replicate, which were plotted individually for each concentration of Didox.

When comparing the two stereoisomers as monotherapies, we found *1R,3R*-RSL3 to be even more potent against *P. falciparum* than *1S,3R*-RSL3 (Figure 6B, Suppl. Fig. 10). While *1R,3R*-RSL3 retained the ability to synergize with Pyrimethamine, the degree of synergy was lower compared to that with *1S,3R*-RSL3 (Figure 6, C and D, Suppl. Fig. 11; cf. Figure 1, C-F).

In summary, both RSL3 stereoisomers exhibit potent antimalarial activity, either as standalone treatments or in combination with Pyrimethamine, albeit with differences in their efficacy and the degree of synergy. These results suggest that specific activity against GPX4 or a related target is not essential for the antiparasitic effect of RSL3.

### Neither thioredoxin reductase inhibitors nor ribonucleotide reductase inhibitors can substitute for RSL3 to synergize with Pyrimethamine

The primary targets of RSL3 responsible for inducing ferroptosis in human cells remain under investigation. While GPX4 [26] was initially described as the main target of RSL3, the compound was later reported to bind and inhibit thioredoxin reductase 1 (TXNRD1) [27]. As a secondary effect, inhibition of TXNRD1 is anticipated to impair the activity of ribonucleotide reductase (RNR), an enzyme crucial for removing the 2’-OH group from ribonucleotides [41]. To evaluate whether these targets contribute to the synergy between RSL3 and Pyrimethamine, we tested additional inhibitors of each enzyme. As thioredoxin reductase inhibitors, we investigated ML162 [27, 42], Auranofin [43] and the TXNRD1 inhibitor 1 (TRi-1) [44]. On the other hand, hydroxyurea [45] and Didox (3,4-dihydroxybenzohydroxamic acid) [46] were used as ribonucleotide reductase inhibitors. Although all these compounds were capable of partially inhibiting the proliferation of *P. falciparum* as monotherapies, none of them demonstrated synergy with Pyrimethamine (Figure 7, A-E, and Suppl. Fig. 12-16). These findings suggest that thioredoxin reductase does not serve as the primary target underlying the synergistic interaction between RSL3 and Pyrimethamine against *P. falciparum*.

### Peroxiredoxin inhibitors (Darapladib, Conoidin A) and other ferroptosis inducers (Erastin, Erastin2, Buthionine sulfoximine) do not synergize with Pyrimethamine against *P. falciparum*

Next, we investigated whether additional inducers of ferroptosis in mammalian cells could synergize with Pyrimethamine to eliminate plasmodia. Erastin and Erastin2 are known antagonists of cellular cystine uptake, reducing the availability of cysteine, which serves as a precursor in glutathione biosynthesis [20]. On the other hand, *L*-Buthionine-*S,R*-sulfoximine (BSO) is an inhibitor of glutamate-cysteine ligase, an enzyme essential for glutathione synthesis [47]. Clinically, BSO is used as a sensitizer in cancer radio-and chemotherapy [48]. In addition to that, BSO was also found to be active against *P. falciparum in vitro* [49]. As a caveat, while all three compounds are predicted to diminish glutathione availability in RBCs, their specific impact on intra-plasmodial glutathione levels remains unclear. In our experiments, Erastin, Erastin2 and BSO inhibited the proliferation of *P. falciparum* as single agents, yet none of them exhibited synergy with Pyrimethamine (Figure 8, A–C, and Suppl. Fig. 17-19).

**Figure 8:**
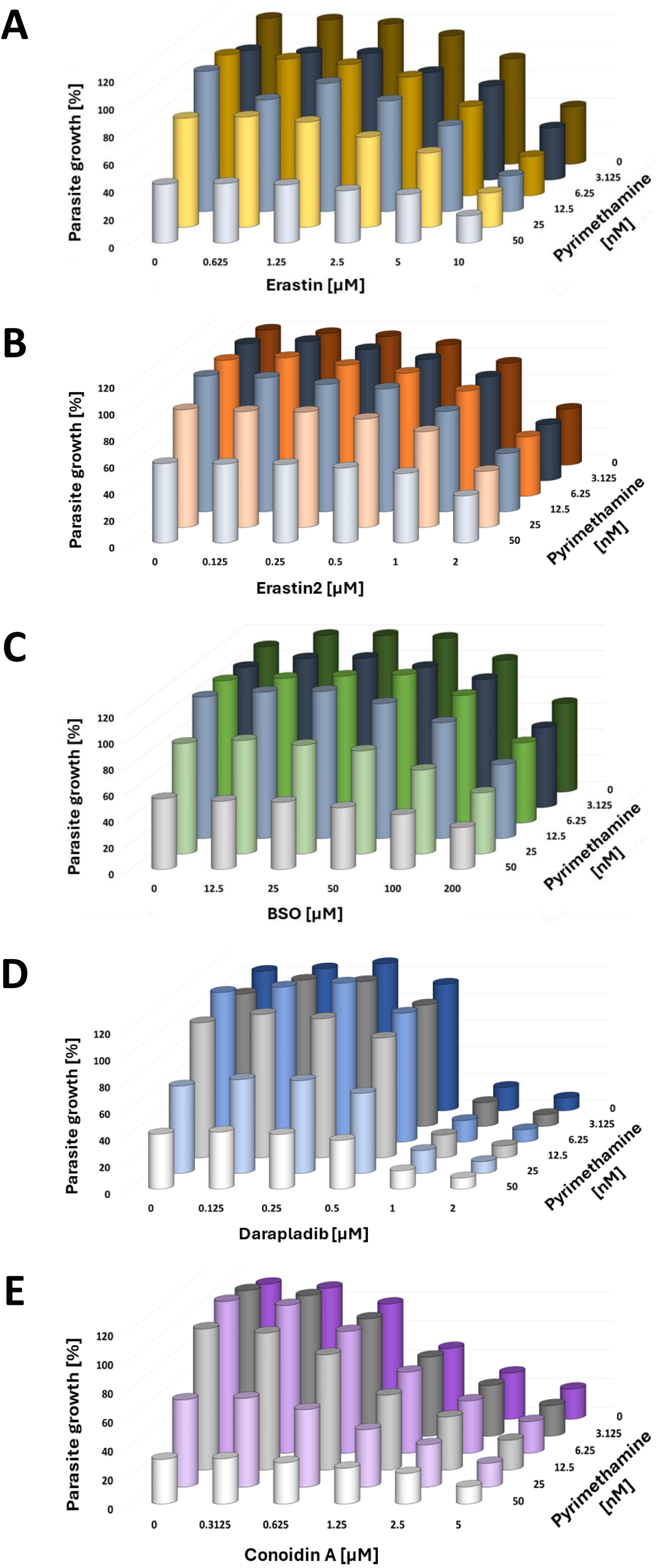
Failure of glutathione antagonists and inhibitors of peroxiredoxins to synergize with Pyrimethamine. **A.** *P. falciparum* parasites were incubated with Erastin (cystine uptake inhibitor) and Pyrimethamine to assess the relative reduction in DNA after 72 hours. No cooperation between the drugs was observed. Suppl. Fig. 17 displays the values obtained from each biological replicate, which were plotted individually for each concentration of Erastin. **B.** Erastin2 (inhibitor of cystine uptake) with Pyrimethamine. Suppl. Fig. 18 displays the values obtained from each biological replicate, which were plotted individually for each concentration of Erastin2. **C.** *L*-Buthionine-*S,R*-sulfoximine (BSO, inhibitor of glutamate-cysteine ligase) with Pyrimethamine. Suppl. Fig. 19 displays the values obtained from each biological replicate, which were plotted individually for each concentration of BSO. **D.** Darapladib (inhibitor of Peroxiredoxin 6) with Pyrimethamine. Suppl. Fig. 20 displays the values obtained from each biological replicate, which were plotted individually for each concentration of Darapladib. **E.** Conoidin A (inhibitor of Peroxiredoxin 2) with Pyrimethamine. Suppl. Fig. 21 displays the values obtained from each biological replicate, which were plotted individually for each concentration of Conoidin A.

**Figure 9:**
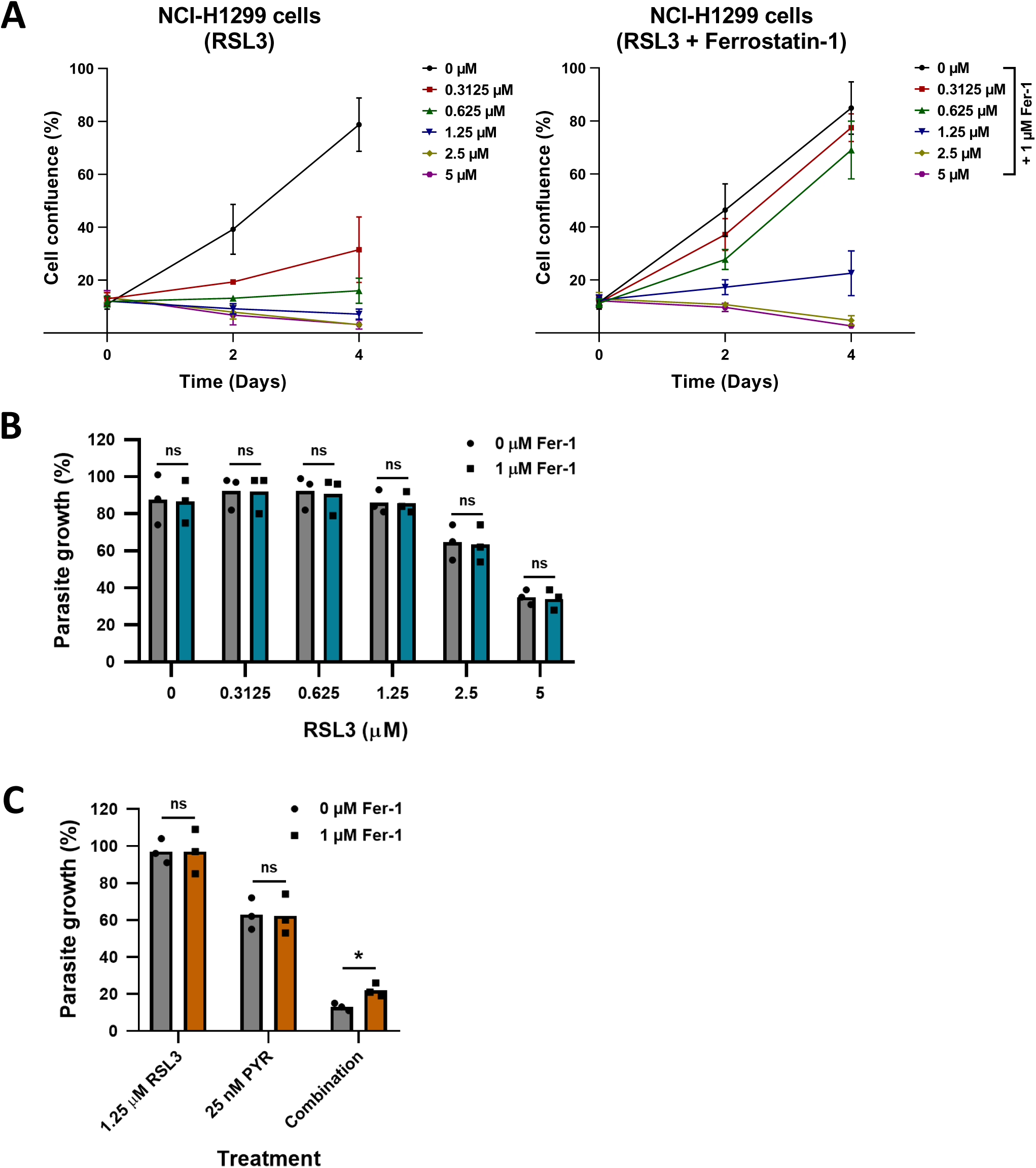
Failure of Ferrostatin-1 to rescue *P. falciparum* from the impact of RSL3. **A.** A widely used human cell line derived from lung adenocarcinoma, NCI-H1299, was incubated with RSL3 and/or Ferrostatin-1. Cell confluence was monitored by automated microscopy to show that Ferrostatin-1 was alleviating the cytotoxicity of RSL3. **B.** Asynchronously growing *P. falciparum* parasites were treated with RSL3, in the presence or absence of Ferrostatin-1. RSL3 suppressed the growth of *P. falciparum* in a dose-dependent manner, regardless of Ferrostatin-1. ns = non-significant **C.** Parasites were treated as in (**B**) but combining the three compounds RSL3, Pyrimethamine, and Ferrostatin-1 as indicated. Ferrostatin-1 provided a small but significant (*, p<0.05) rescue of the parasites from the RSL3/Pyrimethamine combination. ns = non-significant

Similarly, we tested Darapladib, an inhibitor of Peroxiredoxin 6 with known antiparasitic activity [50], and Conoidin A, a permeable inhibitor of Peroxiredoxin 2 exhibiting activity against *P. falciparum* [51]. Although both compounds were effective at blocking the proliferation of *P. falciparum* individually, neither of them showed synergy with Pyrimethamine (Figure 8, D– E, Suppl. Fig. 20 and 21).

These results suggest that none of the commonly employed inducers of ferroptosis or oxidative stress, apart from RSL3, can synergize with Pyrimethamine to effectively eliminate *P. falciparum in vitro*.

### Ferrostatin-1 largely fails to rescue *P. falciparum* from the deleterious impact of RSL3

In the context of ferroptosis, radical-trapping antioxidants (RTAs) are known to rescue cells from the cytotoxic effects of RSL3 [16]. Consistent with this, we observed that Ferrostatin-1, a well-characterized ferroptosis inhibitor and scavenger of lipid hydroperoxides [52], effectively protected human cancer cells from RSL3-induced toxicity. Specifically, preincubation with Ferrostatin-1 rendered NCI-H1299 cells, a human lung cancer cell line, largely resistant to RSL3 treatment, at least when moderate concentrations of RSL3 were used (Figure 9A).

To determine whether a similar rescue effect occurs in *P. falciparum*, we tested Ferrostatin-1 on RSL3-treated parasite cultures. Strikingly, Ferrostatin-1 had no detectable impact on the antiparasitic activity of RSL3 (Figure 9B). Furthermore, when added to the combination of RSL3 and Pyrimethamine, Ferrostatin-1 allowed a significant increase in parasite survival, but it was far from a complete rescue, and most of the parasite growth was still suppressed in the presence of all three compounds (Figure 9C). Within the constraints of the membrane permeability and metabolic stability of Ferrostatin-1 in infected red blood cells, these findings suggest that the mechanism of action of RSL3 against *P. falciparum* might be largely independent of lipid peroxidation.

Finally, we tested the therapeutic efficacy of RSL3 and Pyrimethamine on a different related parasite, *Toxoplasma gondii*. Like *P. falciparum*, *T. gondii* belongs to the phylum *Apicomplexa*. When treating *T. gondii* with either RSL3 or Pyrimethamine, both drugs were inhibitory at low micromolar concentrations. However, combining the drugs did not result in any observable synergy (Suppl. Fig. 22, 23 and 24). This observation points to a species-specific mechanism of action that allows drug synergy in *P. falciparum*.

Taken together, these observations raise the possibility that the mechanism underlying the antiplasmodial effect of RSL3, as well as its synergy with Pyrimethamine and Cycloguanil, is distinct from the well-characterized role of RSL3 in inducing ferroptosis in mammalian cells. Importantly, Ferrostatin-1 or related compounds might be useful when applying RSL3 against malaria in patients, because Ferrostatin-1 would ameliorate the toxicity of RSL3 towards human cells without substantially compromising its therapeutic efficacy against plasmodia, thus providing a useful therapeutic window.

## DISCUSSION

In this study, we demonstrate that RSL3, a compound originally developed to induce ferroptosis in mammalian cells, exhibits synergistic activity with the established antimalarial drugs Pyrimethamine and Cycloguanil to eliminate *P. falciparum* within human RBCs. This synergy highlights the potential of RSL3 or its derivatives as candidates for repurposing in malaria treatment, particularly in combination with diamino compounds like Pyrimethamine.

Despite the promising drug synergy observed, the underlying target structures within *P. falciparum* and/or their host RBCs remain unclear. This is particularly true for Pyrimethamine and its structurally related analogue Cycloguanil. Both drugs have *P. falciparum* DHFR as their only well-characterized target among *Plasmodium* spp., a notion supported by the frequent mutations in the *DHFR* gene observed in Pyrimethamine-resistant parasites [8, 15, 35, 53, 54]. However, other established *P. falciparum* DHFR inhibitors did not demonstrate synergy with RSL3 in our study. This suggests that Pyrimethamine may target additional sites, and that the synergy observed with RSL3 likely results from the inhibition of at least one of these alternative targets.

The target responsible for the observed synergy may involve an alternative activity of *P. falciparum* DHFR, which is targeted by Pyrimethamine but not by Methotrexate or WR99210. One possibility is that the thymidylate synthase (TS) activity, which resides within the same DHFR/TS fusion protein in *P. falciparum* [55], could be involved. Another intriguing possibility comes from recent findings in humans, according to which DHFR catalyses the synthesis of tetrahydrobiopterin (BH_4_), and failure to do so increases cellular sensitivity to RSL3 (Soula et al., 2020). If a similar activity exists for *P. falciparum* DHFR, and if Pyrimethamine and Cycloguanil interfere with it, while other DHFR inhibitors do not, this could explain why the synergy with RSL3 is limited to these two drugs. In support of this idea, RSL3 has been reported to synergize with Methotrexate to eliminate human cancer cells, where Methotrexate inhibits BH_4_ regeneration [56, 57]. Further investigation is needed to determine whether *P. falciparum* DHFR can also regenerate BH_4_, whether this confers resistance to RSL3, and whether this activity of *P. falciparum* DHFR is inhibited by Pyrimethamine and Cycloguanil but not by Methotrexate or WR99210.

Additional targets of Pyrimethamine within *P. falciparum* or their host RBCs could also help explain the synergy with RSL3. Notably, Pyrimethamine has been described as an inhibitor of the transcription factor STAT3 in human cells [58] and as a mediator that destabilizes aminoacyl-tRNA synthetase-interacting multifunctional protein 2 (AIMP2) [59]. Other potential targets include NF-κB, MAPK, thymidine phosphorylase and telomerase [60]. Although many of these targets are not present in *P. falciparum*, these findings suggest that Pyrimethamine could exert effects on a broader range of targets in *P. falciparum*, beyond just DHFR. This widens the potential for establishing effective synergies with other broadly active drugs like RSL3.

The activity of RSL3 against *P. falciparum* and its synergy with Pyrimethamine were unique among the pro-ferroptotic compounds we studied. Other inducers of ferroptosis did not replace RSL3 to synergize with Pyrimethamine. Most strikingly, Ferrostatin-1, an antagonist of lipid peroxidation, did not rescue *P. falciparum* from the impact of RSL3 and had only weak rescue activity against the combination of RSL3 and Pyrimethamine. These observations suggest that RSL3 may cooperate with Pyrimethamine and act against *P. falciparum* in a manner largely independent of oxidative stress or ferroptosis-related reactions.

Additionally, RSL3 may have a variety of target proteins in both human cells and protozoa. Previous work suggested that GPX4 [26, 61] or thioredoxin reductase [27] could be targeted by RSL3. However, more specific inhibitors of thioredoxin reductase or ribonucleotide reductase failed to synergize with Pyrimethamine against *P. falciparum* in our assays. Recent pre-published work suggested selenoproteins as potential targets of RSL3 [28]. These proteins are encoded by a stop codon in mRNA and synthesized through the incorporation of selenocysteine, after modifying tRNA-bound serine [62]. Previous studies have identified at least four selenoproteins in *P. falciparum* [63, 64]. Thus, in principle, RSL3 could exert its action by targeting and modifying any of these selenoproteins in *P. falciparum*. However, we cannot exclude the possibility that RSL3 also targets proteins of the host RBCs. The chloroacetamide moiety of RSL3 may form covalent bonds with selenohydryl groups, as well as sulfhydryl or imidazole groups [23, 24], suggesting a wide range of potential targets in RBCs or parasites that could interfere with the proliferation of plasmodia.

**Limitations of the study**: It is important to note that the drug synergy was observed *in vitro*, i.e. in cultured *P. falciparum* inside RBCs. Whether this synergy holds true in an *in vivo* model, such as *P. berghei* in mice, remains to be determined. Additionally, for potential clinical translation, the pharmacokinetics and toxicology of RSL3 need to be investigated, as it has not yet undergone published clinical trials. Of note, however, anti-ferroptotic molecules like Ferrostatin-1 can alleviate the toxicity of RSL3 on human cells but not on *P. falciparum*, which might open a therapeutic window. Another limitation is the potential impact of Pyrimethamine-resistant *P. falciparum* on the efficacy of the combination therapy with RSL3. However, it is worth mentioning that not all *P. falciparum* DHFR mutations result in complete resistance [9, 14], and Pyrimethamine derivatives have been shown to regain activity against some of these DHFR mutants [65].

**Perspectives**: With RSL3 or related compounds in hand, it may be possible to enhance therapies targeting not just the intraerythrocytic but also the liver stages of *Plasmodium* infections. For instance, combining RSL3 with Proguanil, which is metabolized to Cycloguanil, could offer a promising strategy. This approach could be beneficial not only for preventing *P. falciparum* infections but also for eliminating chronic infections caused by *Plasmodium vivax*. While the lack of well-defined RSL3 targets in *P. falciparum* represents a challenge, it also offers an exciting opportunity for future research and development of novel therapeutic strategies. Our study suggests that drugs containing chloroacetamide or similar reactive moieties, capable of covalently modifying their target proteins, could be effective against malaria parasites. This opens up the potential for large-scale screening efforts to identify such modifiers, similar to previous screens conducted in human cells [23, 66, 67], which could lead to the discovery of new antimalarial compounds and uncover previously “undruggable” targets within plasmodia.

## ACKNOWLEDGEMENTS

During this work, VN was a scholar of the International Max Planck Research School for Molecular Biology, part of the GGNB graduate school.

The authors are grateful to the Infectious Diseases Imaging Platform (www.idip-heidelberg.org) and the Plasmodium database PlasmoDB (www.plasmodb.org), which facilitated this work. This study was supported by the Health + Life Science Alliance Heidelberg Mannheim, receiving state funding approved by the State Parliament of Baden-Württemberg, and the German Research Foundation - project number 240245660 - SFB 1129 to MG.

## AUTHOR CONTRIBUTIONS

Conceptualization: MD

Experimentation: VN, KL, AZ, CGL

Supervision: MG, MD

Manuscript, initial draft: MD

Manuscript, correction and finalization: all authors

## MATERIALS AND METHODS

### Cell culture

The human non-small cell lung cancer cell line NCI-H1299 was maintained in Dulbecco’s Modified Eagle’s Medium (DMEM, Thermo Fisher Scientific), which was supplemented with 10% Fetal Bovine Serum (FBS, Anprotec), 2 mM glutamine (Thermo Fisher Scientific), 50 U/mL penicillin, 50 μg/mL streptomycin (Gibco), 10 μg/mL ciprofloxacin (Fresenius Kabi) and 2 μg/mL tetracycline (Sigma-Aldrich). Cells were cultivated at 37°C at 5% CO_2_. They were routinely tested for *Mycoplasma* contamination and scored negative.

### Confluence measurements of human NCI-H1299 cells

NCI-H1299 cells were seeded on transparent 12-well plates (Corning Costar) and continuously treated with the indicated compounds. The treatment was started on the day after seeding, and the cells were treated for a total of 4 days, during which the treatment was refreshed every other day. Cell confluence was measured using an automated imaging cytometer (Celigo, Nexcelom Bioscience). For data analysis and graph generation, GraphPad Prism 8.0 was used.

### Cultivation of *P. falciparum* 3D7 in human RBCs

*P. falciparum*, strain 3D7, was obtained from Hosam Shams-Eldin (Marburg University, Germany). The asexual blood stages were cultivated in AB-positive or AB-negative human RBCs (German Red Cross blood bank, DRK-NSTOB) in homemade RPMI-1640 medium (Gibco) supplemented with 5.95 g/L HEPES (Carl Roth), 50 mg/L hypoxanthine (Carl Roth), 0.2% NaHCO_3_ (Carl Roth), 0.5% AlbuMAX-I (Gibco) and 12.5 µg/mL gentamycin (Sigma Aldrich). RBCs were maintained at approx. 4% hematocrit. Parasites were maintained at 37°C in hypoxia incubator chambers (STEMCELLTechnologies) under a hypoxic atmosphere consisting of 95% N_2_, 5% CO_2_ and 5% O_2_.

### Drug treatment, calculation of IC_50_ values and determination of synergy scores

Both RSL3 isomers (*1S,3R*-RSL3 and *1R,3R*-RSL3), Erastin, Erastin2, Buthionine sulfoximine, Cycloguanil hydrochloride and WR99210 were obtained from Hycultec (MedChemExpress). Dihydroartemisinin, Atovaquone, Pyrimethamine, Darapladib, were provided by Selleckchem. All compounds were reconstituted in DMSO, and were used at the indicated concentrations in the respective assays. For IC_50_ value calculation, nonlinear regression (curve fit) in GraphPad Prism was used. Bliss synergy scores were calculated using the *Synergy Finder* software provided by www.synergyfinder.org.

### Giemsa staining of thin blood smears and light microscopy for assessment of parasite morphology

Thin blood smears were made on glass microscope slides (Labsolute). Next, parasites were fixed in methanol for 1-5 minutes. Smears were allowed to dry completely and then incubated with Giemsa’s solution (Sigma Aldrich) diluted 1:4 in Giemsa buffer [11.7 µM Na_2_HPO_4_ (Carl Roth), 4 µM KH_2_PO_4_ (Carl Roth) in ddH_2_O] for 10 minutes at room temperature. After the staining, slides were rinsed under running water and dried. A Zeiss Axio Scope.A1 microscope equipped with a Zeiss Achroplan 100x/1.25 oil immersion objective was used for quantification and assessment of parasite morphology. Images were taken and processed using ZEN 2 (blue edition).

### Synchronization of *P. falciparum* 3D7

Parasites were synchronized by lysis of late stages using 5% D-sorbitol (Sigma Aldrich). Non-infected and *P. falciparum*-infected RBCs were centrifuged, followed by a 15-minute incubation with 5% D-sorbitol at 37°C. Following that, parasites were washed with complete RPMI-1640 medium by several rounds of centrifugation. Alternatively, parasites were synchronized by MACS^®^ cell separation using LD magnetic columns attached to a MidiMACS™ Separator (Miltenyi Biotec), following the recommendations of the manufacturer. This method takes advantage of the paramagnetic properties of late-stage parasites, allowing separation of early and late parasite stages.

### SYBR Green I fluorescence-based proliferation assay for determination of total DNA content in *P. falciparum*-infected RBCs

Total DNA content of *P. falciparum-*infected RBCs was determined by the SYBR Green I fluorescence-based proliferation assay, with minor modifications of the original protocol [68]. Briefly, 0.5% mixed-stage parasites were incubated with the compounds of interest in black 96-well plates with a translucent bottom (Falcon) for 72 hours under a hypoxic atmosphere. At the end of the treatment, parasites were lysed using homemade lysis buffer [ddH_2_O, 1x SYBR Green I (Invitrogen), 40 mM Tris-HCl pH 7.5 (Carl Roth), 10 mM EDTA pH 8.0 (Carl Roth), 0.02% Saponin (Carl Roth), 0.08% Triton X-100 (Sigma Aldrich)]. DNA levels were measured using a Twinkle LB970 fluorescence plate reader (Berthold Technologies) at 485 nm/520 nm as excitation/emission wavelengths, respectively. Microsoft Excel was used for data analysis and to generate the 3D bar charts.

### Colorimetric assay for determination of *P. falciparum* LDH activity

For determination of *P. falciparum* LDH activity, parasites were synchronized using 5% D-sorbitol, following the procedure described above. After synchronization, late-stage parasites at 1% parasitemia and 2% hematocrit were treated with the compounds of interest for 48 hours. An NBT solution was prepared by adding a nitroblue tetrazolium tablet (NBT, Sigma Aldrich) to a homemade LDH assay buffer [ddH_2_O, 100 mM Tris-HCl pH 8.0 (Carl Roth), 0.2M sodium L-lactate (Sigma Aldrich), 0.25% Triton X-100 (Sigma Aldrich)]. The complete LDH substrate solution, containing the NBT solution, 50 units/ml diaphorase from *Clostridium kluyveri* (Sigma Aldrich) and 10 mg/ml 3-Acetylpyridine adenine dinucleotide (APAD, Sigma Aldrich) was added to the parasites. Parasites were incubated at room temperature for 30-60 minutes. Absorbance was then measured at 650 nm using an Apollo LB913 microplate spectrophotometer (Berthold Technologies). Data analysis was conducted in Microsoft Excel, and GraphPad Prism was used to generate the bar charts.

### *Toxoplasma gondii* and host cell infection

African green monkey kidney epithelial (Vero) cells were cultivated in DMEM (Life Technologies Corporation), supplemented with 10% fetal bovine serum (FBS, Sigma-Aldrich), 2 mM L-glutamine (Biochrom), 100 U/mL penicillin and 100 µg/mL streptomycin (Life Technologies Corporation) at 37°C in a humidified air atmosphere enriched to 5% CO_2_. Cells were infected with mouse-virulent *Toxoplasma gondii* (*T. gondii*) type I reporter strains RH-pSAG-βGal, kindly provided by F. Seeber, Berlin [69] or RH-pTUB-tdTomato RFP, kindly provided by M. Blume, Berlin [70]. *T. gondii* tachyzoites were propagated in L929 fibroblasts as host cells. For infection of Vero cells, freshly egressed parasites were separated from host cells by differential centrifugation following a previously described protocol [71]. After extensive washing, parasites were added to confluent Vero cells in 96-well tissue culture plates at a multiplicity of infection (MOI) of 0.5 to 1. Cells were treated with the indicated concentrations of Pyrimethamine and RSL3 or combinations thereof at 30 minutes post infection (p.i.). At 72 hours p.i., β-galactosidase activity of RH-pSAG-βGal parasites was measured using chlorophenol red-β-D-galactopyranoside (CPRG) as a substrate. To this end, cells were fixed by adding an equal volume of 4% formaldehyde, 250 mM NaCl, 24 mM NaOH in Hank’s Balanced Salt Solution (HBSS; Gibco/Thermo Fisher Scientific). After 20 minutes, cells were washed with HBSS for 15 minutes, followed by incubation with 1 mM CPRG (Sigma-Aldrich), 0.1% Triton X-100 and 5 mM MgCl_2_ in HBSS for 1 hour at 37°C. Absorbance was measured at 570 nm using a BioTek Epoch2 microplate reader (Agilent, Santa Clara, USA). Fluorescence as a measure of RH-pTUB-tdTomato RFP growth was determined at 138 hours p.i. in a Wallac Victor^3^V multilabel counter (PerkinElmer, Waltham, USA) using 544/15 nm excitation and 580/10 nm emission filters.

## LEGENDS TO SUPPLEMENTAL FIGURES

**Supplemental Figure 1:**
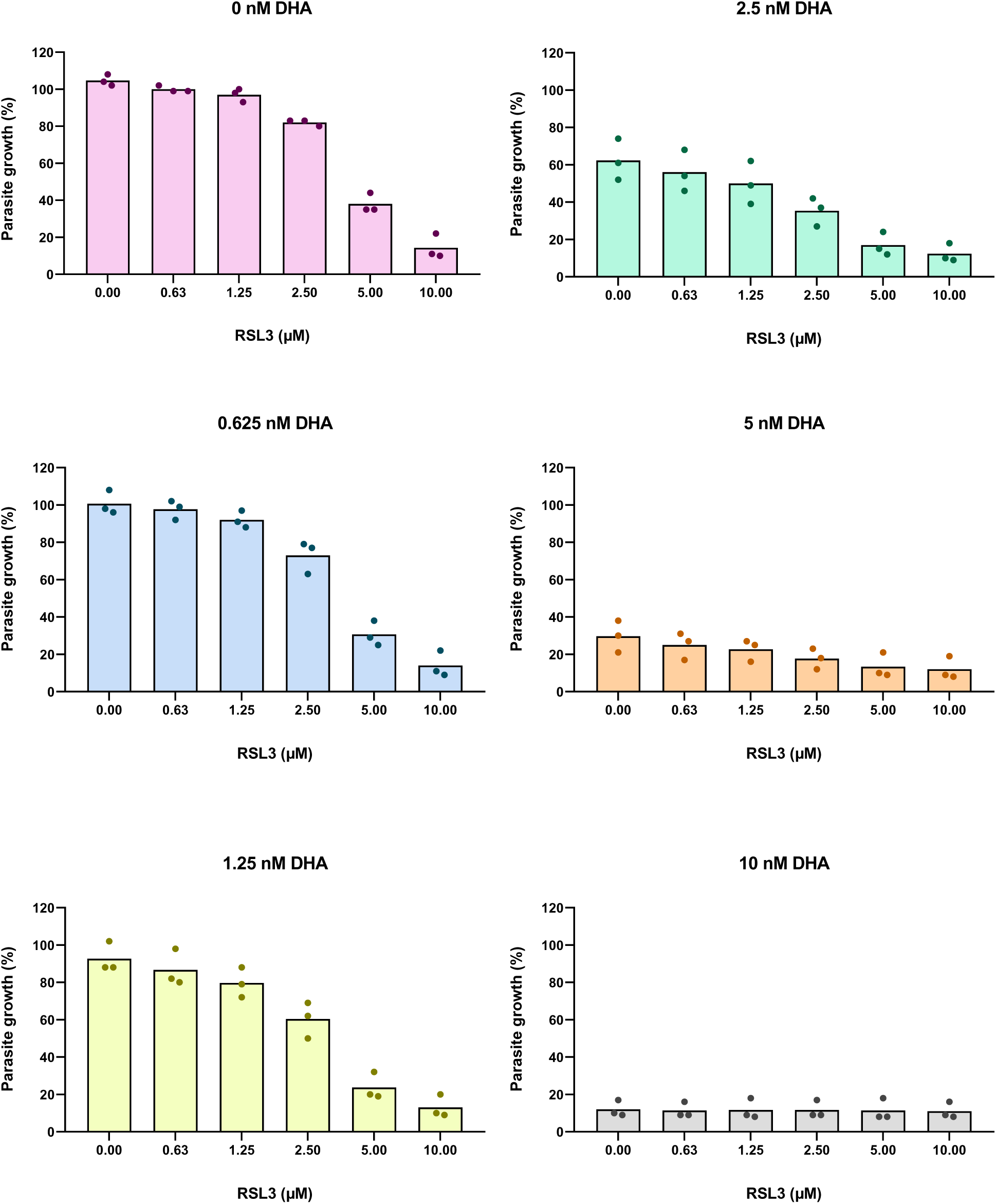
Asynchronously growing *P. falciparum* parasites were treated with RSL3 and Dihydroartemisinin (DHA) at the indicated concentrations. After 72 hours, parasite growth was assessed by measurement of the total DNA content (SYBR Green staining). The values obtained from each biological replicate were plotted individually for each concentration of DHA. Corresponding to Figure 1A

**Supplemental Figure 2:**
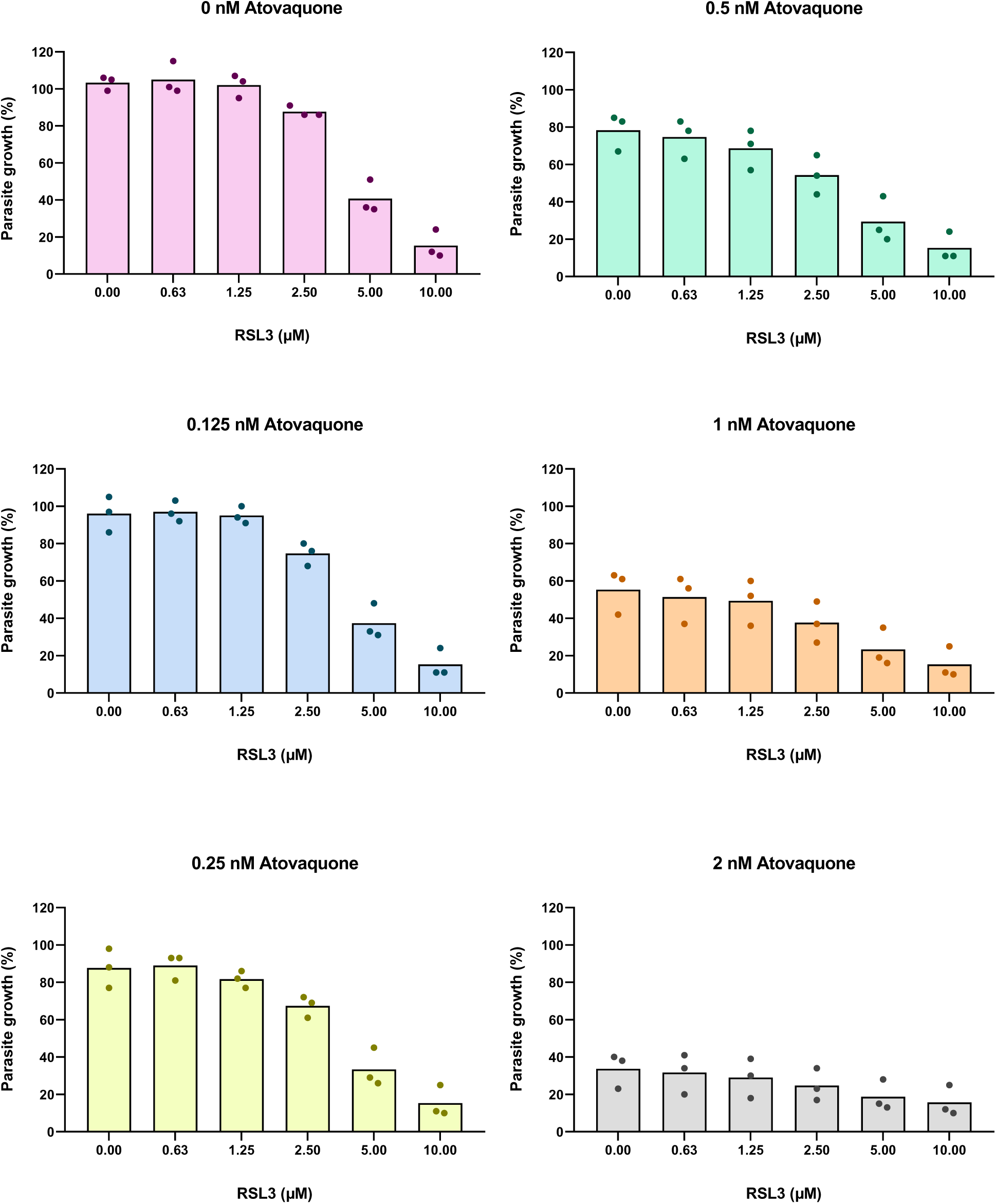
Treatment with RSL3 and Atovaquone as described in Suppl. Fig. 1. Corresponding to Figure 1B

**Supplemental Figure 3:**
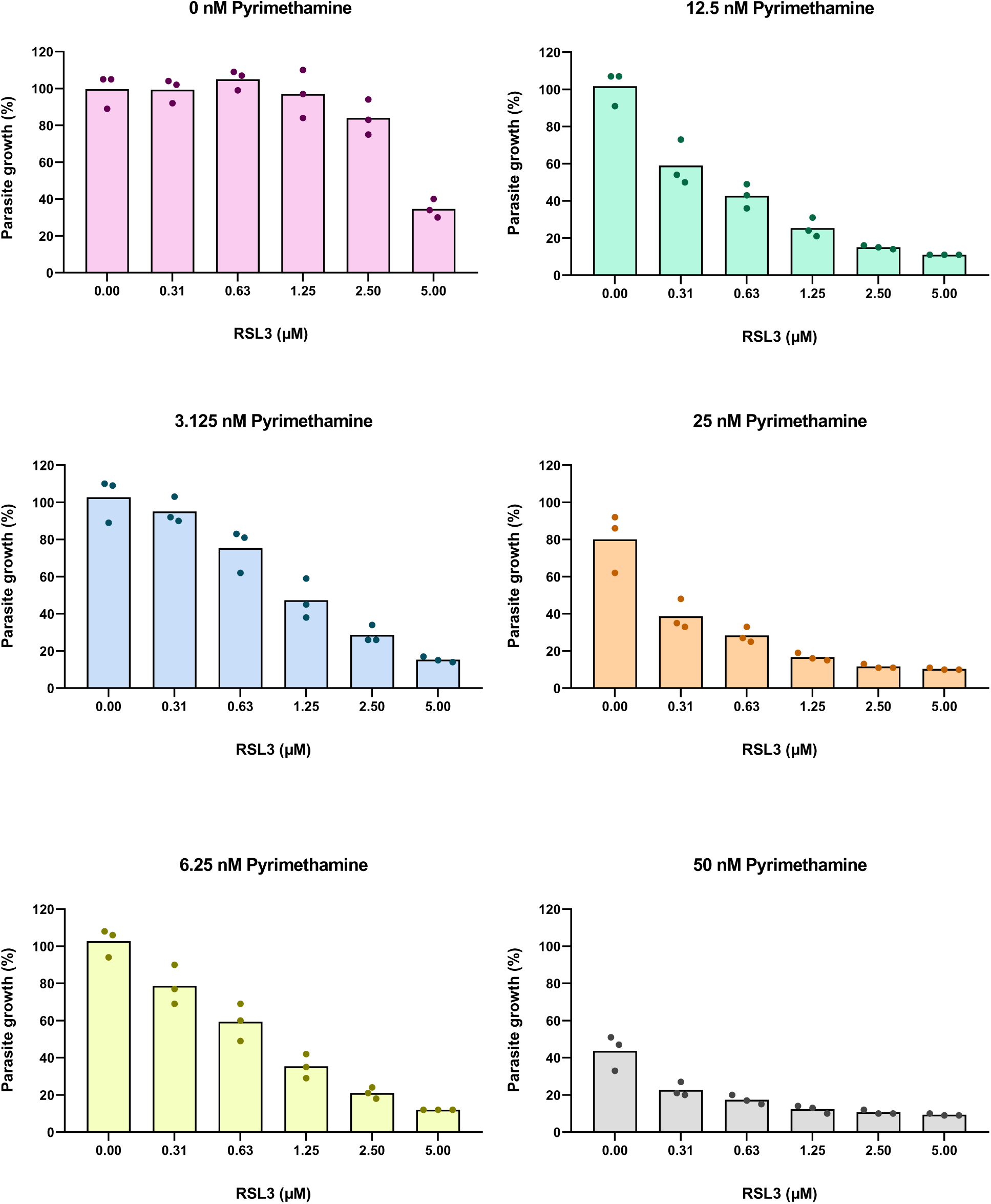
Treatment with RSL3 and Pyrimethamine as described in Suppl. Fig. 1. Corresponding to Figure 1C

**Supplemental Figure 4:**
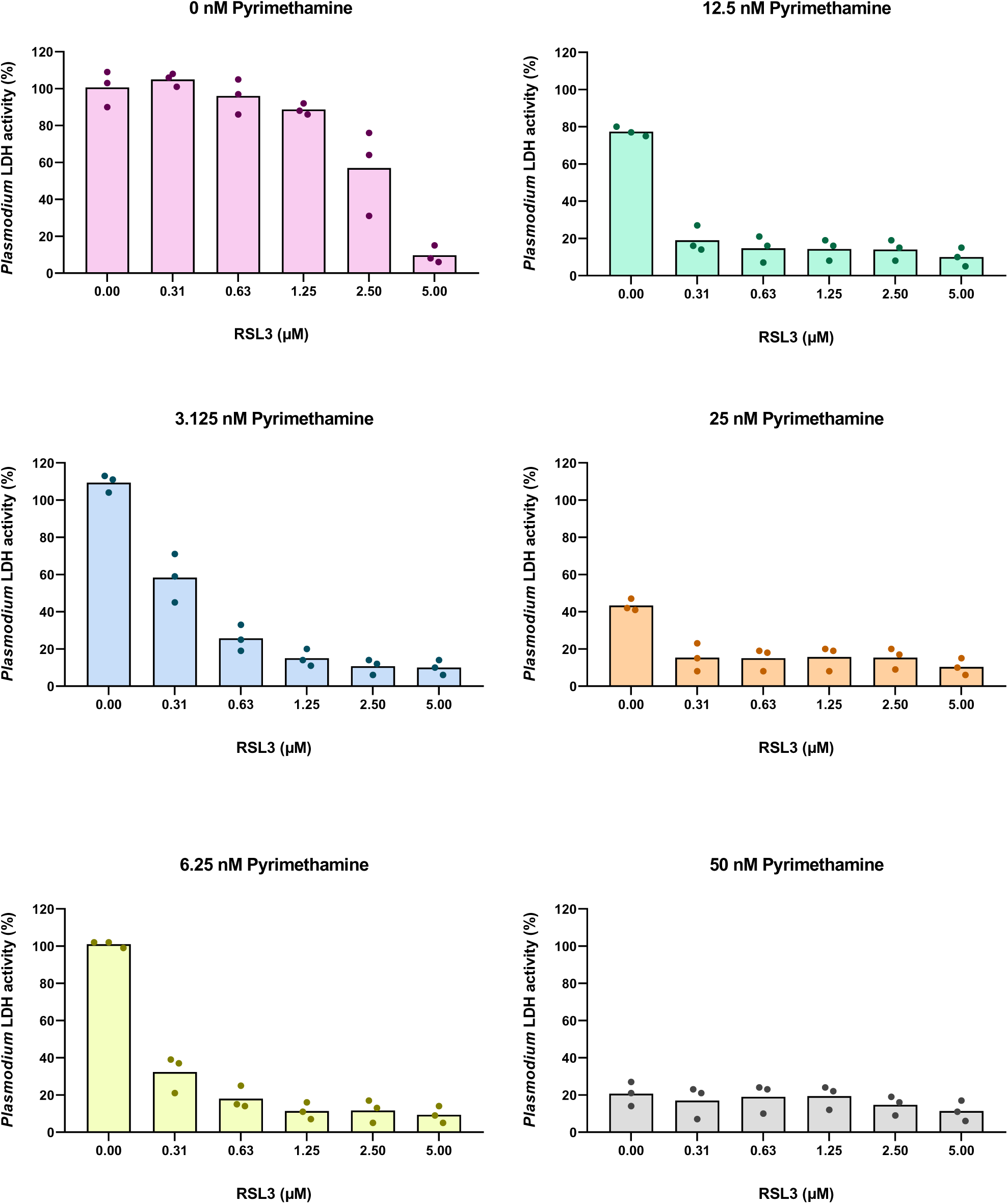
*P. falciparum* parasites were synchronized by sorbitol treatment. Late-stage parasites were treated with RSL3 and Pyrimethamine at the indicated concentrations for 48 hours. As a readout for parasite growth, the activity of *P. falciparum* LDH was measured. The values obtained from each biological replicate were plotted individually for each concentration of Pyrimethamine. Corresponding to Figure 1E

**Supplemental Figure 5:**
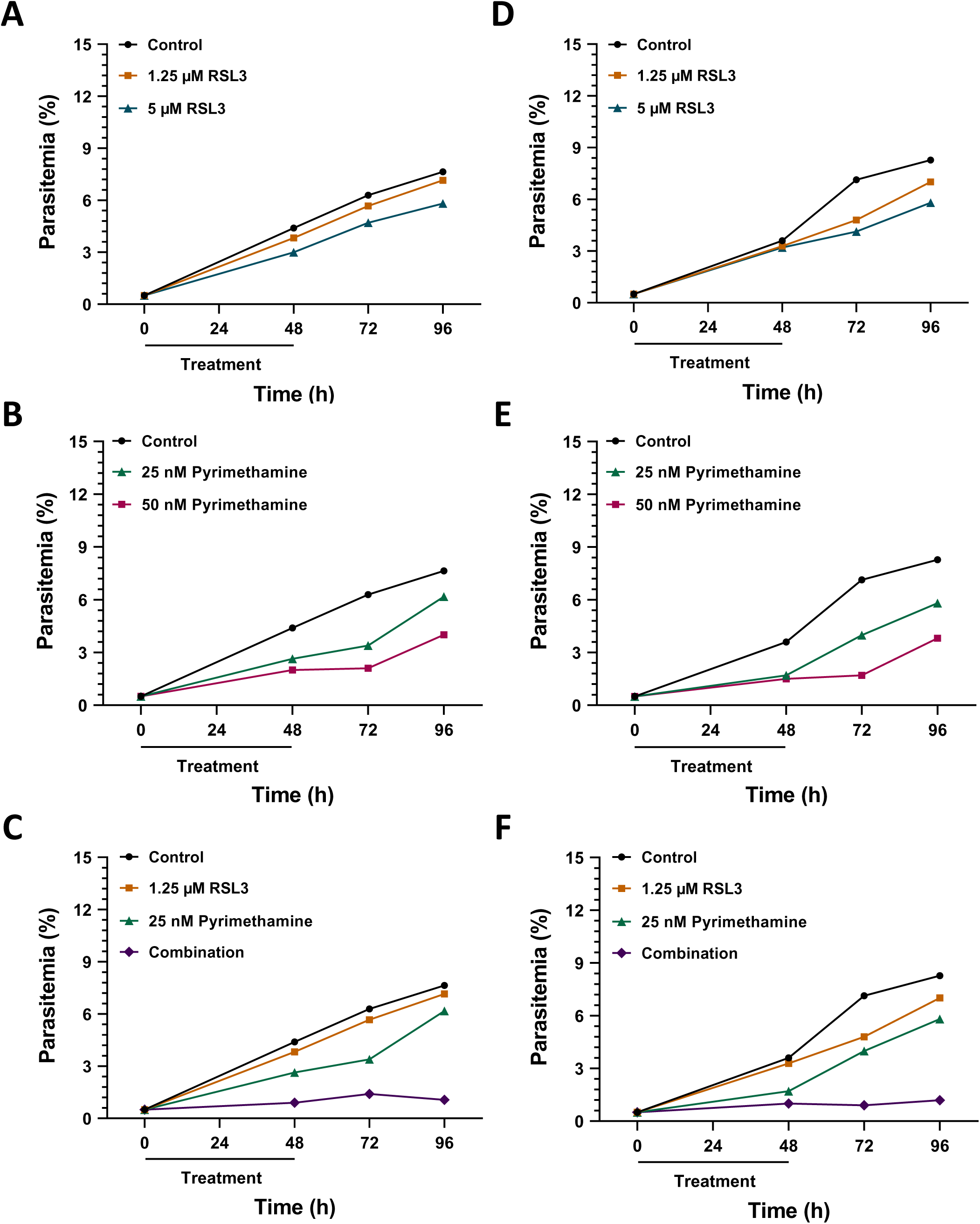
Sustainable inhibition of *P. falciparum* growth by RSL3 and Pyrimethamine, second and third biological replicate. Corresponding to Figure 2 **A.** *P. falciparum* parasites were treated with RSL3 as in Figure 2A (second replicate). **B.** Parasites were exposed to Pyrimethamine as in Figure 2B (second replicate). **C.** Treatment with RSL3 and Pyrimethamine as in Figure 2C (second replicate). **D.** *P. falciparum* parasites were treated with RSL3 as in Figure 2A (third replicate). **E.** Parasites were exposed to Pyrimethamine as in Figure 2B (third replicate). **F.** Treatment with RSL3 and Pyrimethamine as in Figure 2C (third replicate).

**Supplemental Figure 6:**
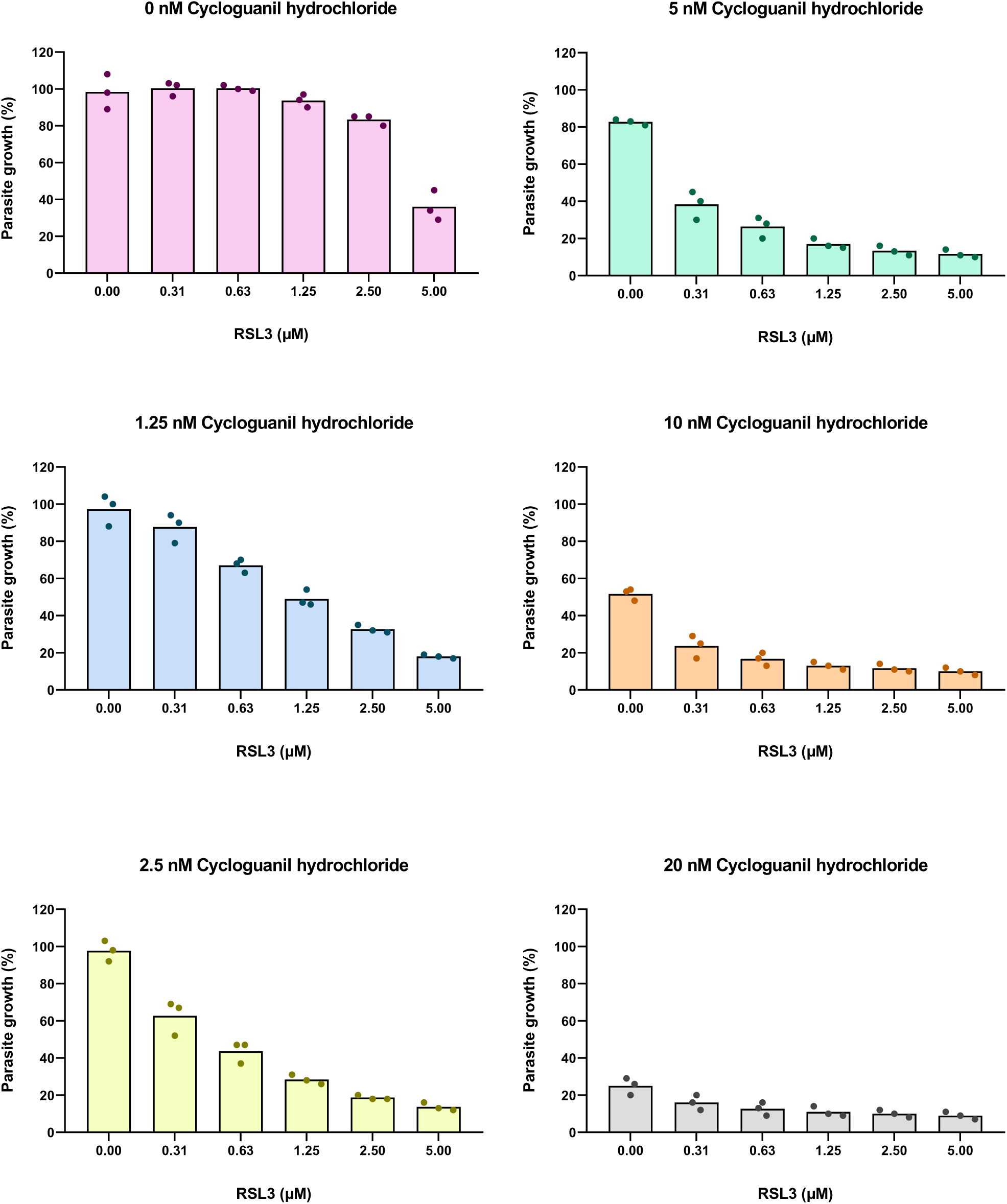
Treatment with RSL3 and Cycloguanil hydrochloride as described in Suppl. Fig. 1. Corresponding to Figure 4B

**Supplemental Figure 7:**
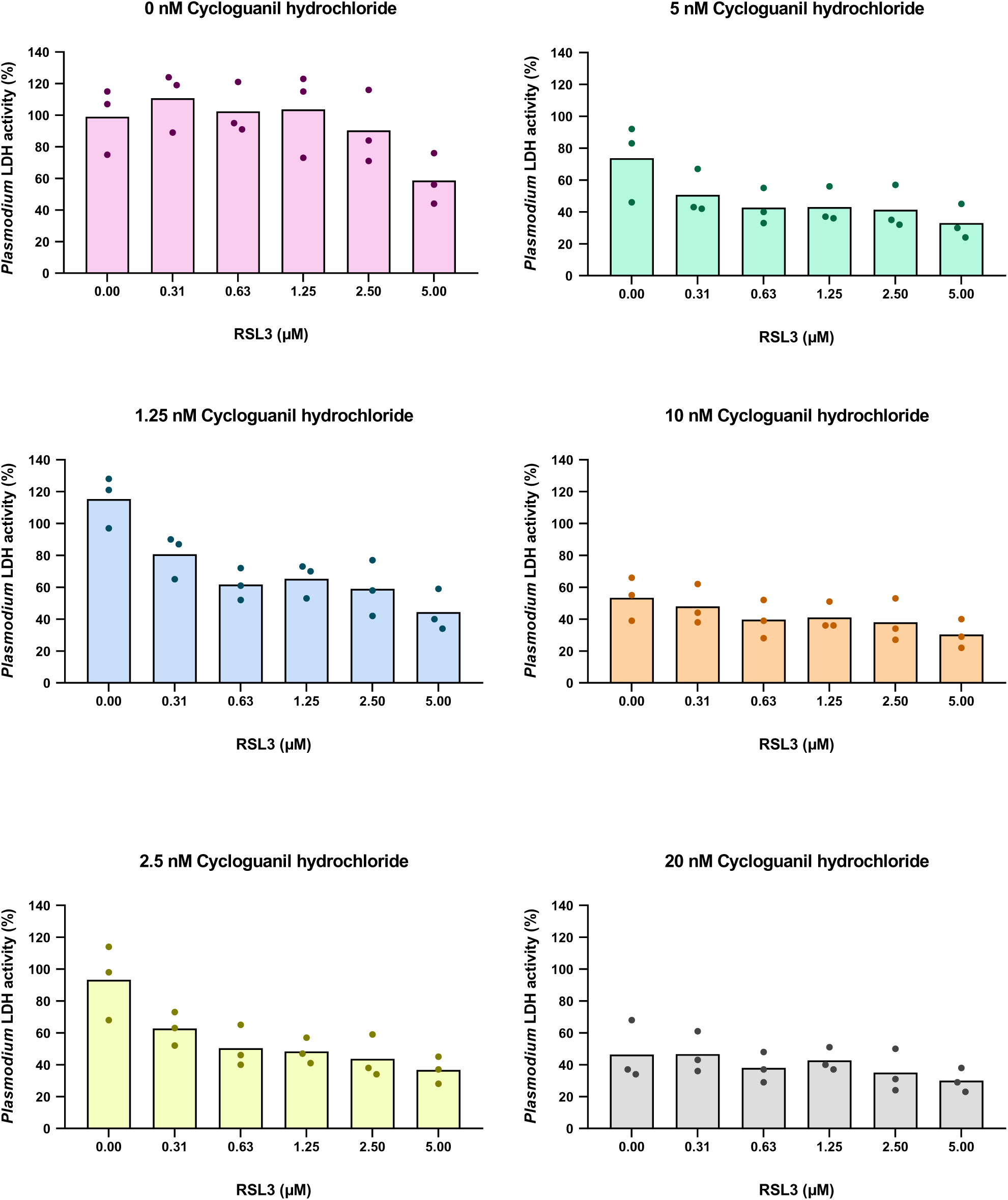
Treatment with RSL3 and Cycloguanil hydrochloride as described in Suppl. Fig. 4. Corresponding to Figure 4D

**Supplemental Figure 8:**
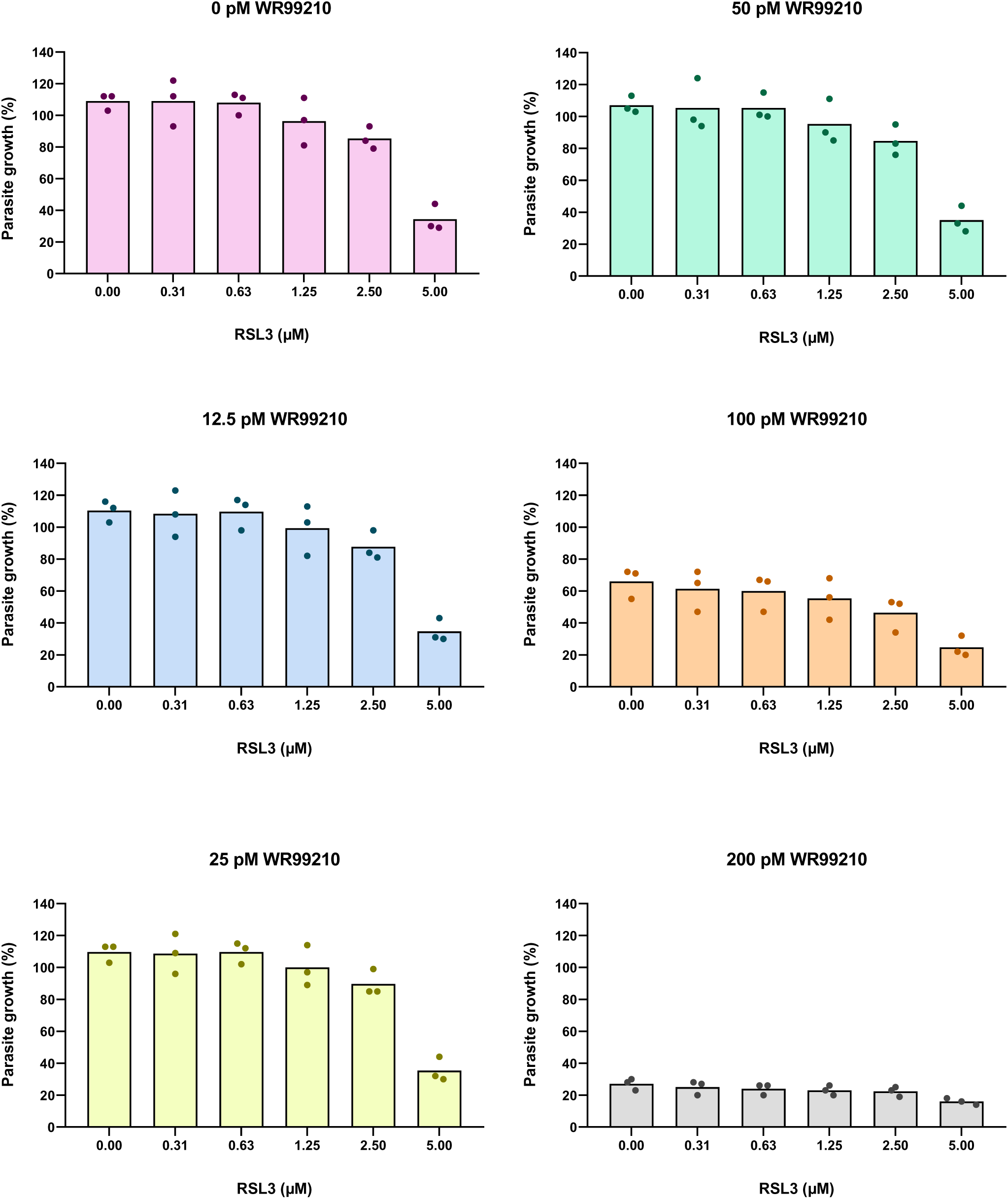
Treatment with RSL3 and WR99210 as described in Suppl. Fig. 1. Corresponding to Figure 5A

**Supplemental Figure 9:**
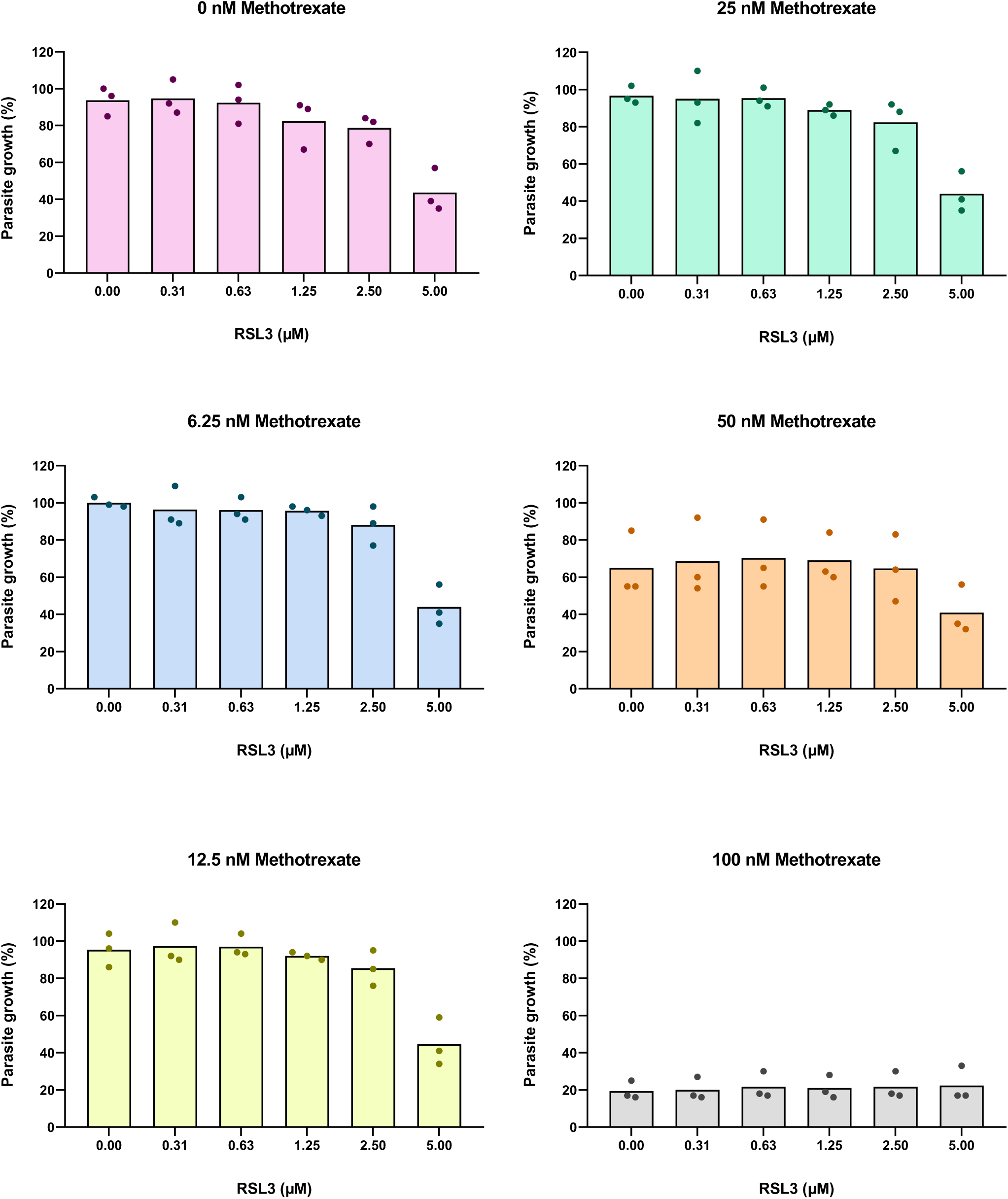
Treatment with RSL3 and Methotrexate as described in Suppl. Fig. 1. Corresponding to Figure 5B

**Supplemental Figure 10:**
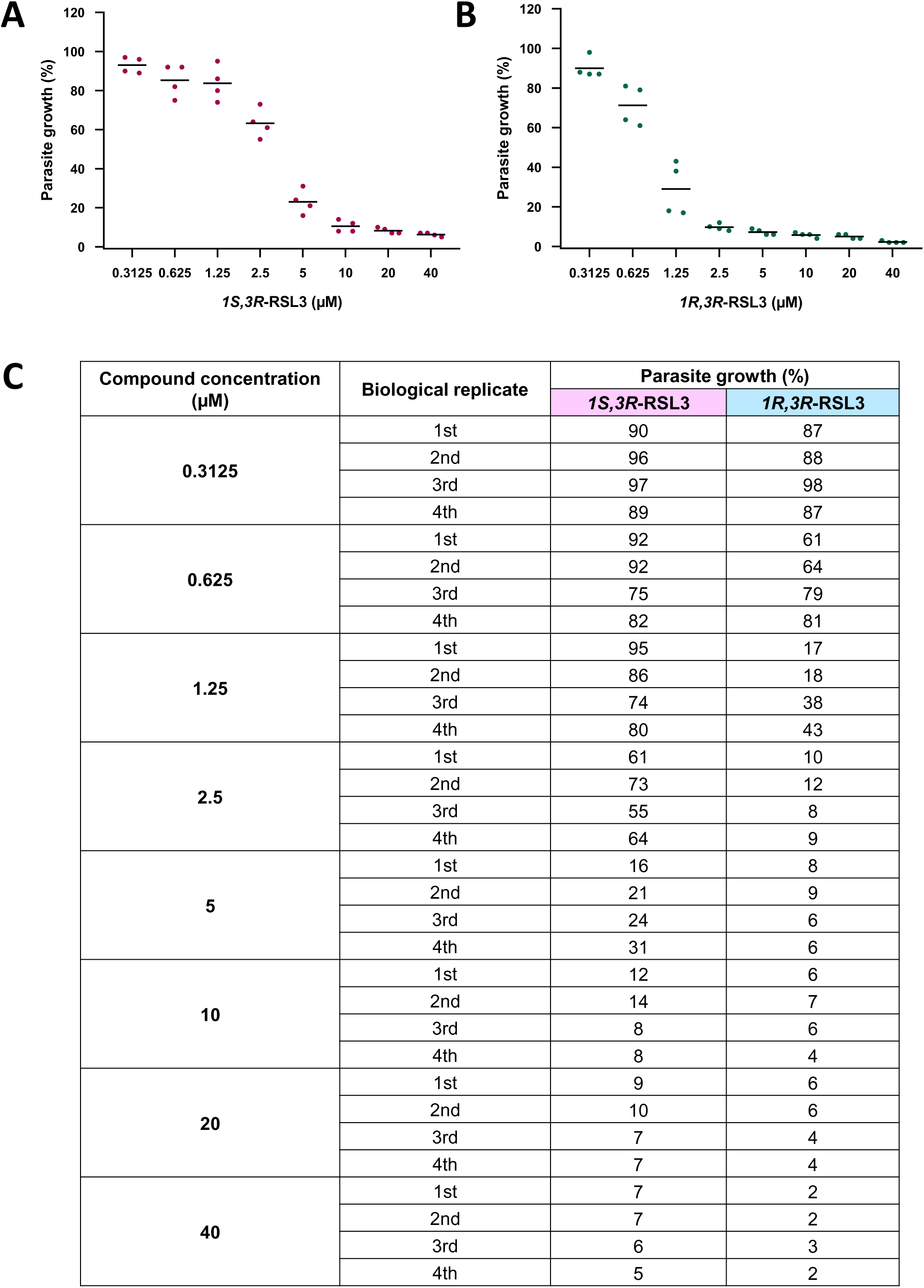
Impact of *1S,3R*-RSL3 and *1R,3R*-RSL3 on the intraerythrocytic proliferation of *P. falciparum*. Corresponding to Figure 6B **A.** Parasite growth in response to the indicated concentrations of *1S,3R*-RSL3 was assessed by total DNA content measurement using the SYBR Green I fluorescencebased proliferation assay. The values obtained from each biological replicate were plotted. **B.** *P. falciparum* parasites were treated with *1R,3R*-RSL3 and their proliferation was evaluated as described in (**A**). **C.** Quantification of parasite growth (percentage) from three biological replicates for both *1S,3R*-RSL3 and *1R,3R*-RSL3.

**Supplemental Figure 11:**
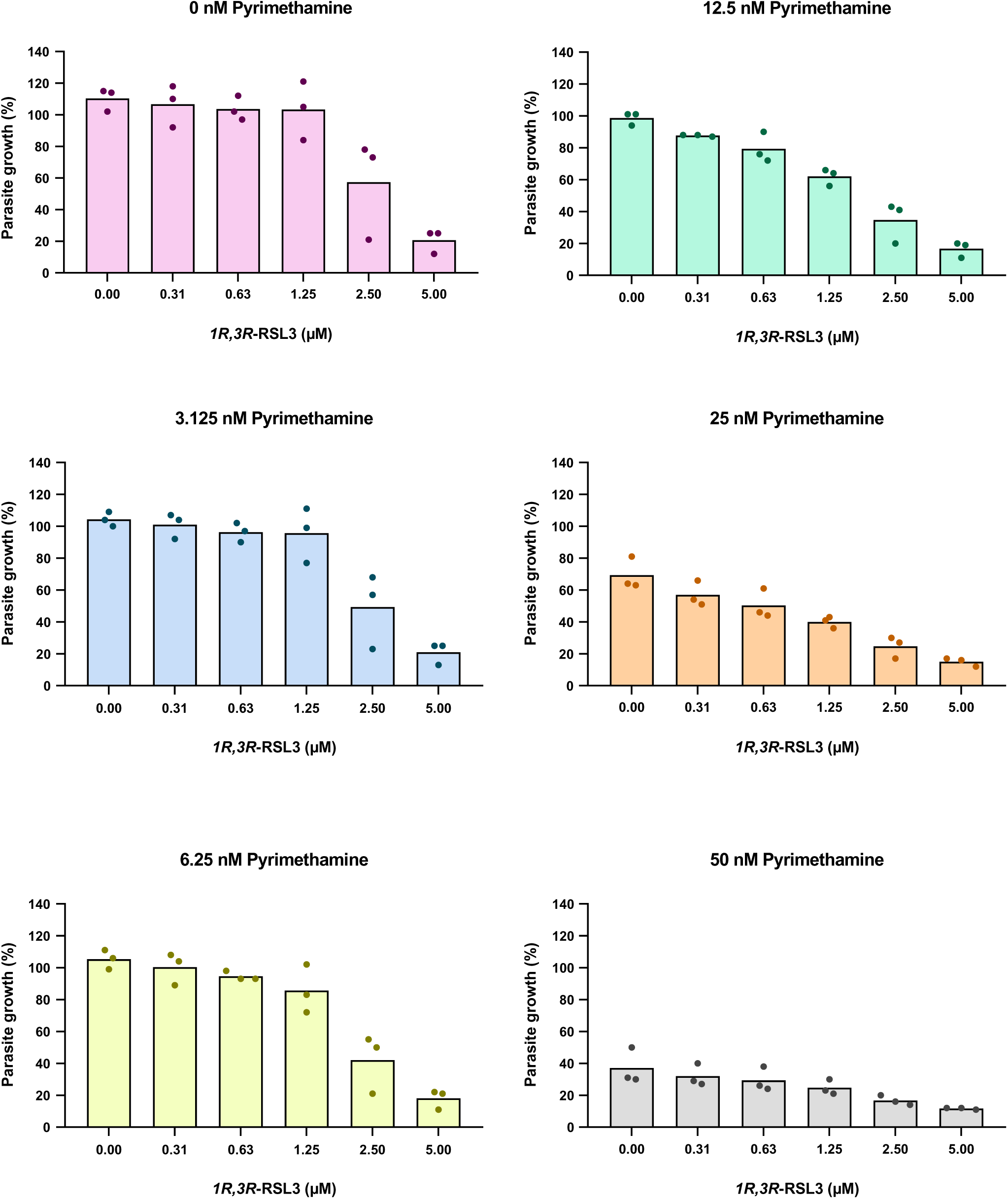
Treatment with *1R,3R*-RSL3 and Pyrimethamine as described in Suppl. Fig. 1. Corresponding to Figure 6C

**Supplemental Figure 12:**
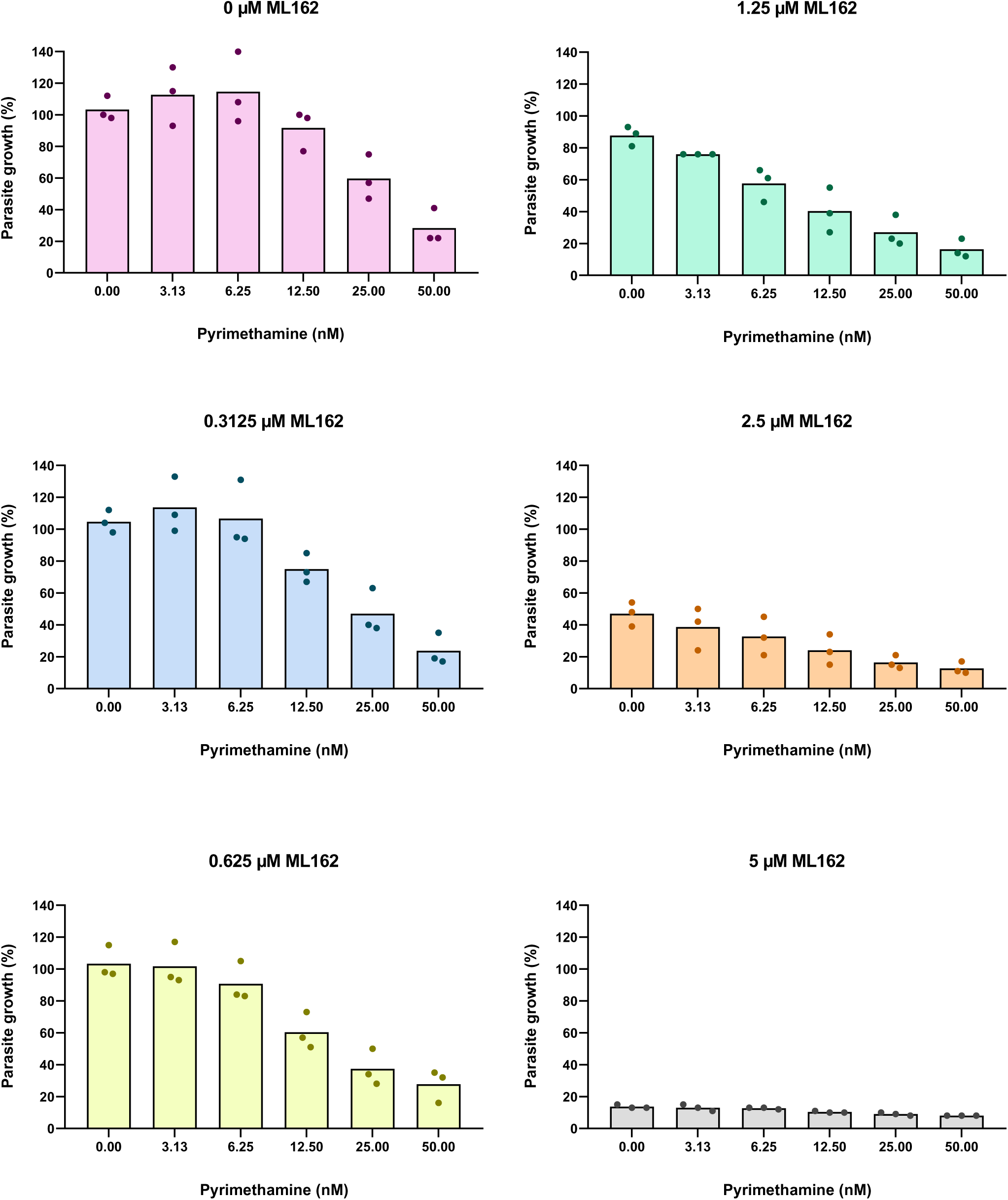
Treatment with Pyrimethamine and ML162 as described in Suppl. Fig. 1. Corresponding to Figure 7A

**Supplemental Figure 13:**
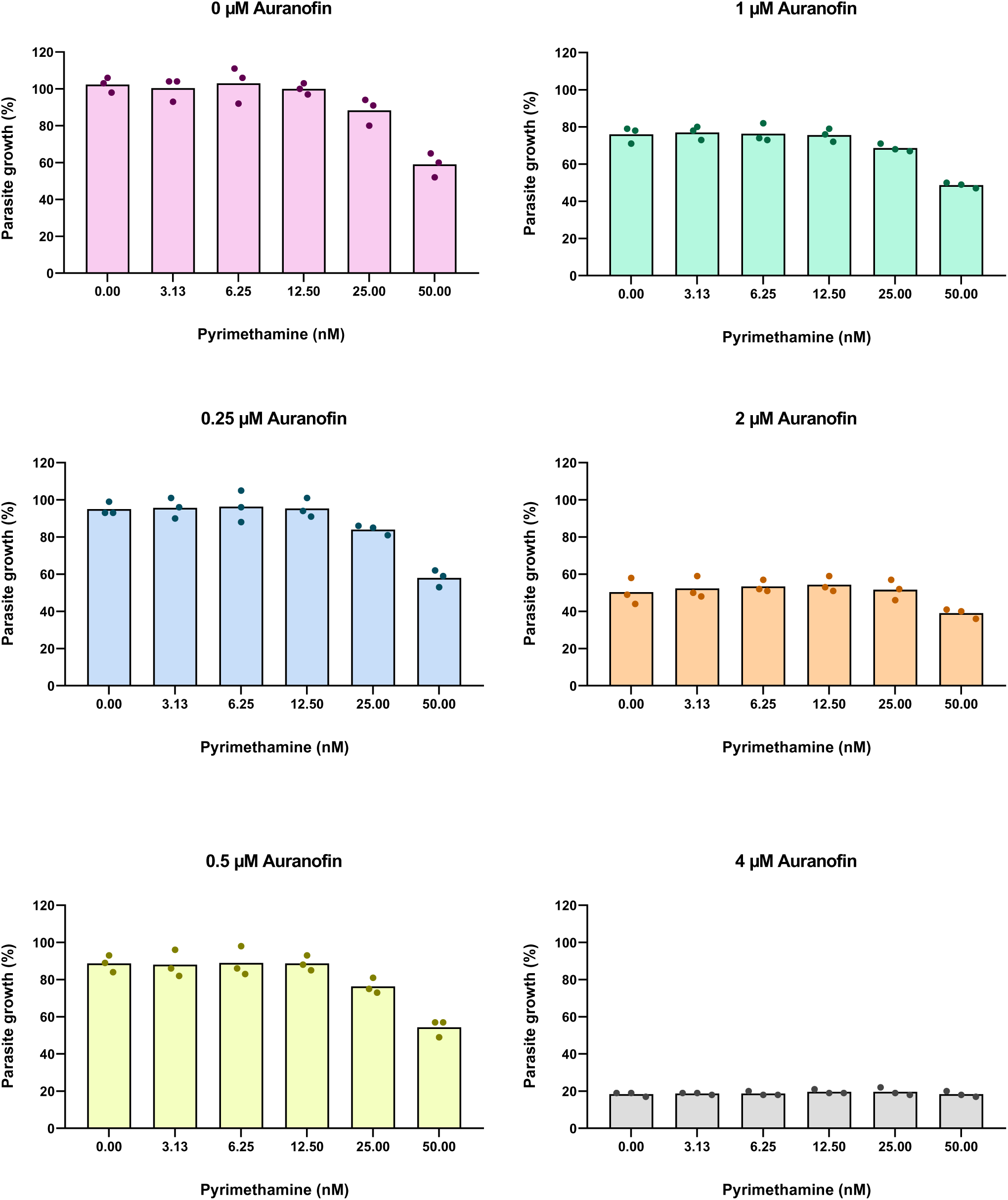
Treatment with Pyrimethamine and Auranofin as described in Suppl. Fig. 1. Corresponding to Figure 7B

**Supplemental Figure 14:**
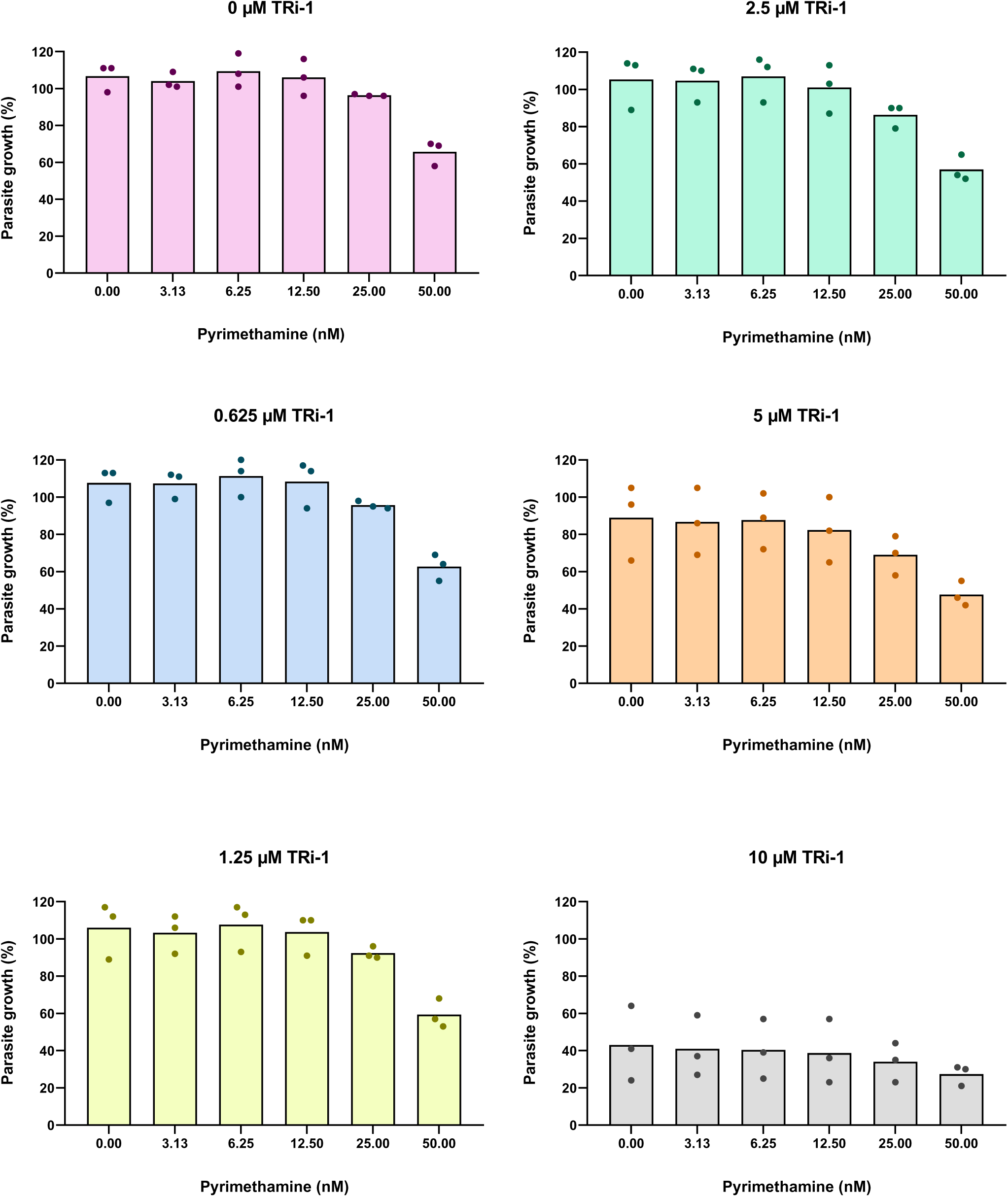
Treatment with Pyrimethamine and TRi-1 as described in Suppl. Fig. 1. Corresponding to Figure 7C

**Supplemental Figure 15:**
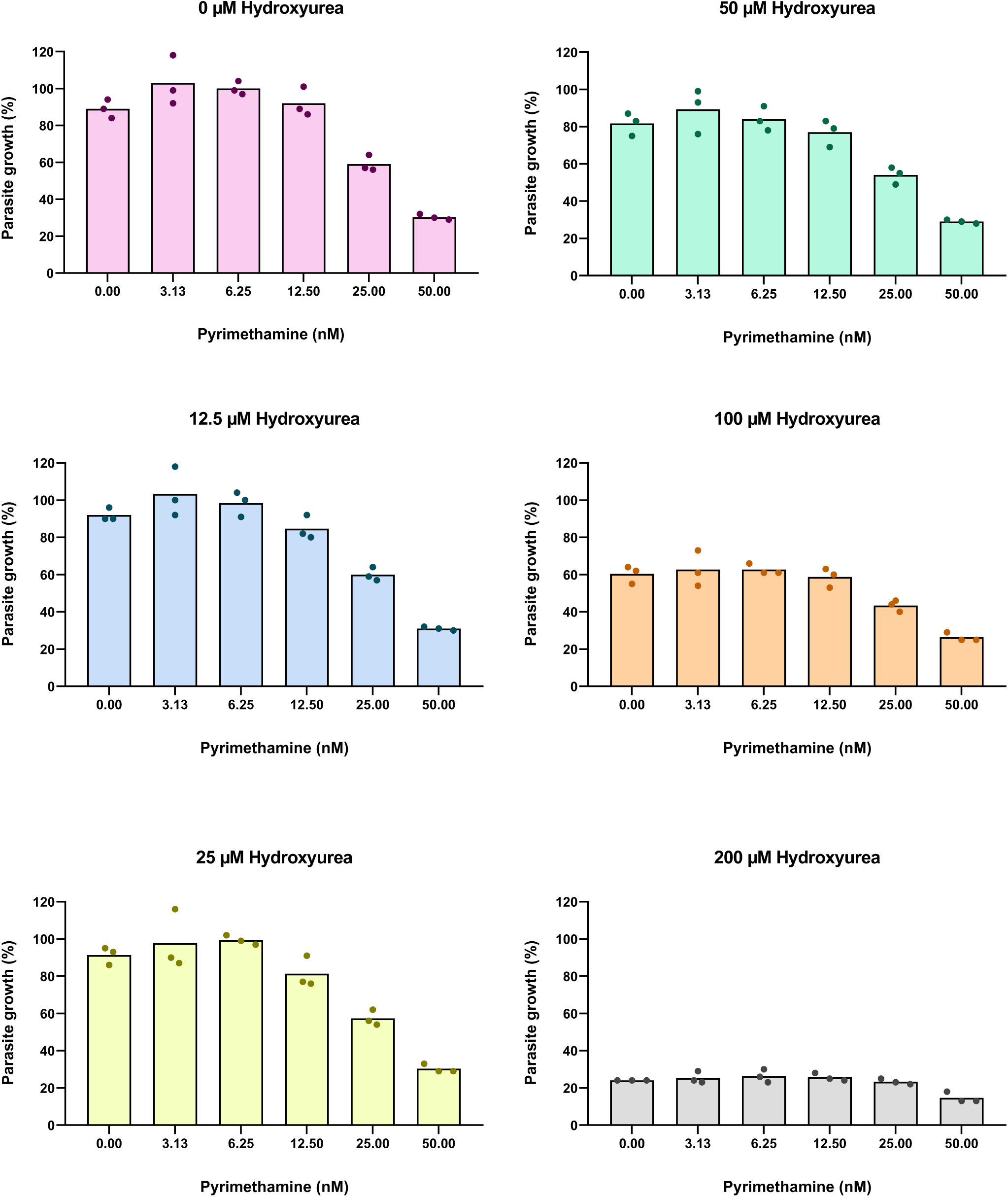
Treatment with Pyrimethamine and Hydroxyurea as described in Suppl. Fig. 1. Corresponding to Figure 7D

**Supplemental Figure 16:**
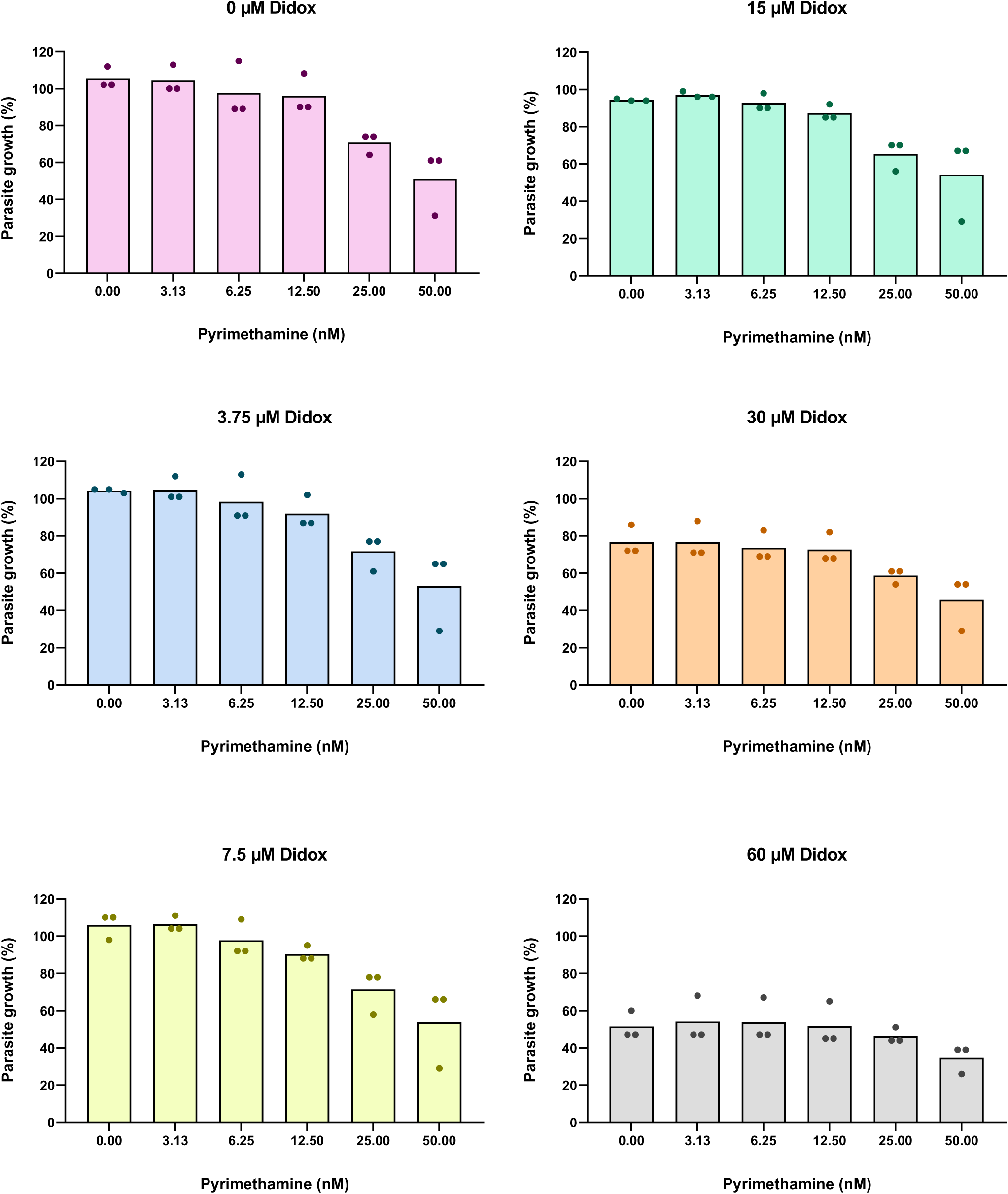
Treatment with Pyrimethamine and Didox as described in Suppl. Fig. 1. Corresponding to Figure 7E

**Supplemental Figure 17:**
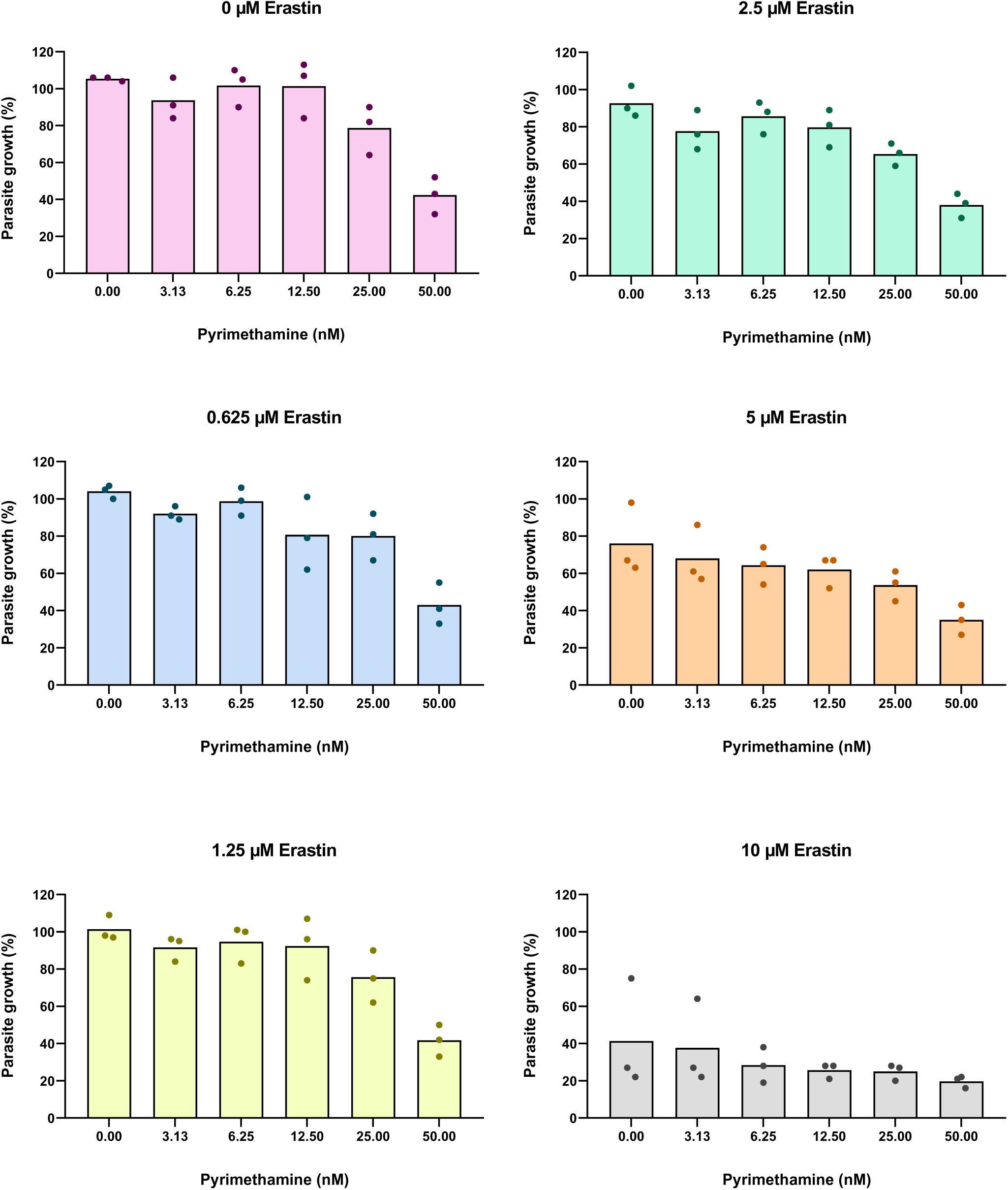
Treatment with Pyrimethamine and Erastin as described in Suppl. Fig. 1. Corresponding to Figure 8A

**Supplemental Figure 18:**
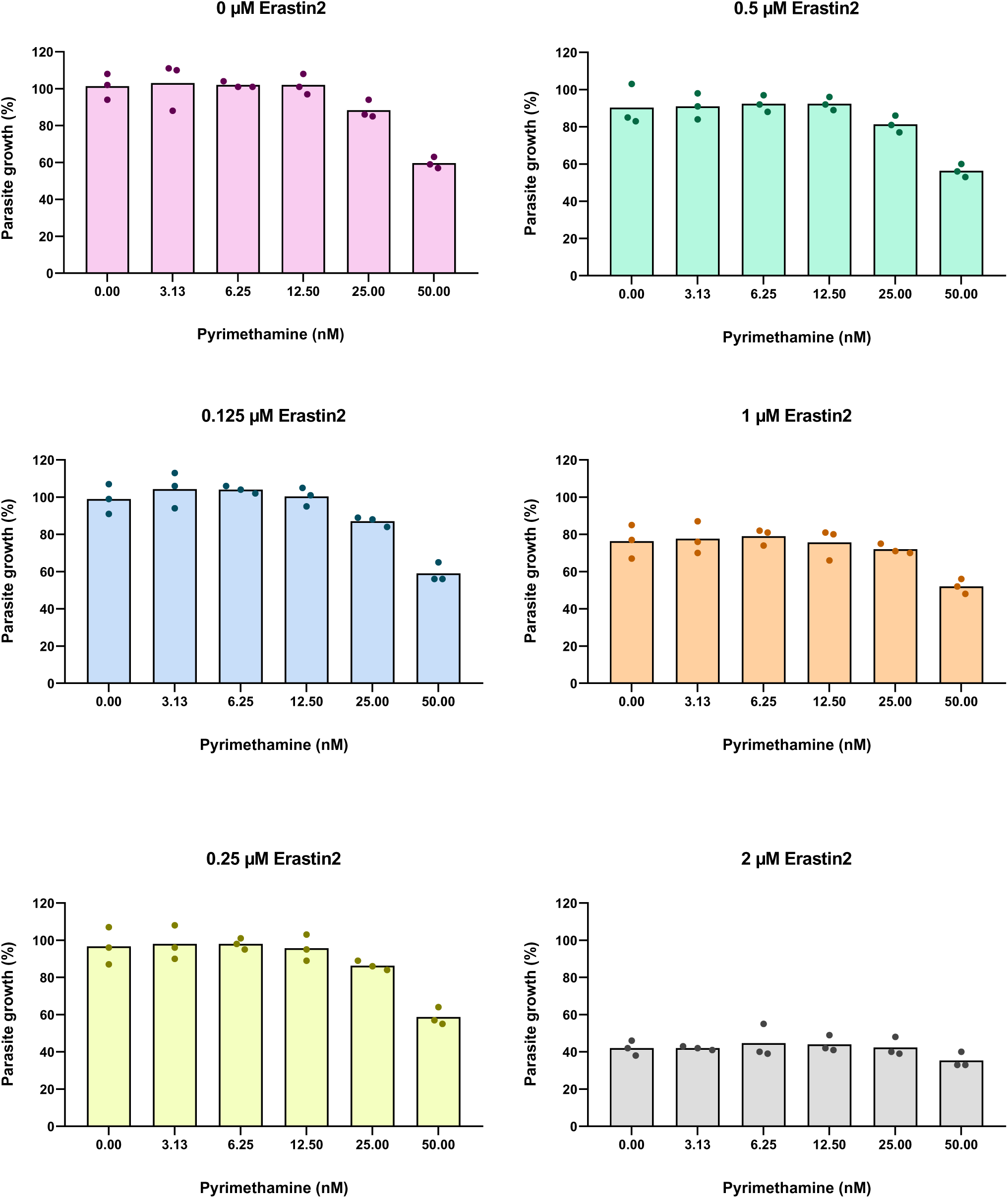
Treatment with Pyrimethamine and Erastin2 as described in Suppl. Fig. 1. Corresponding to Figure 8B

**Supplemental Figure 19:**
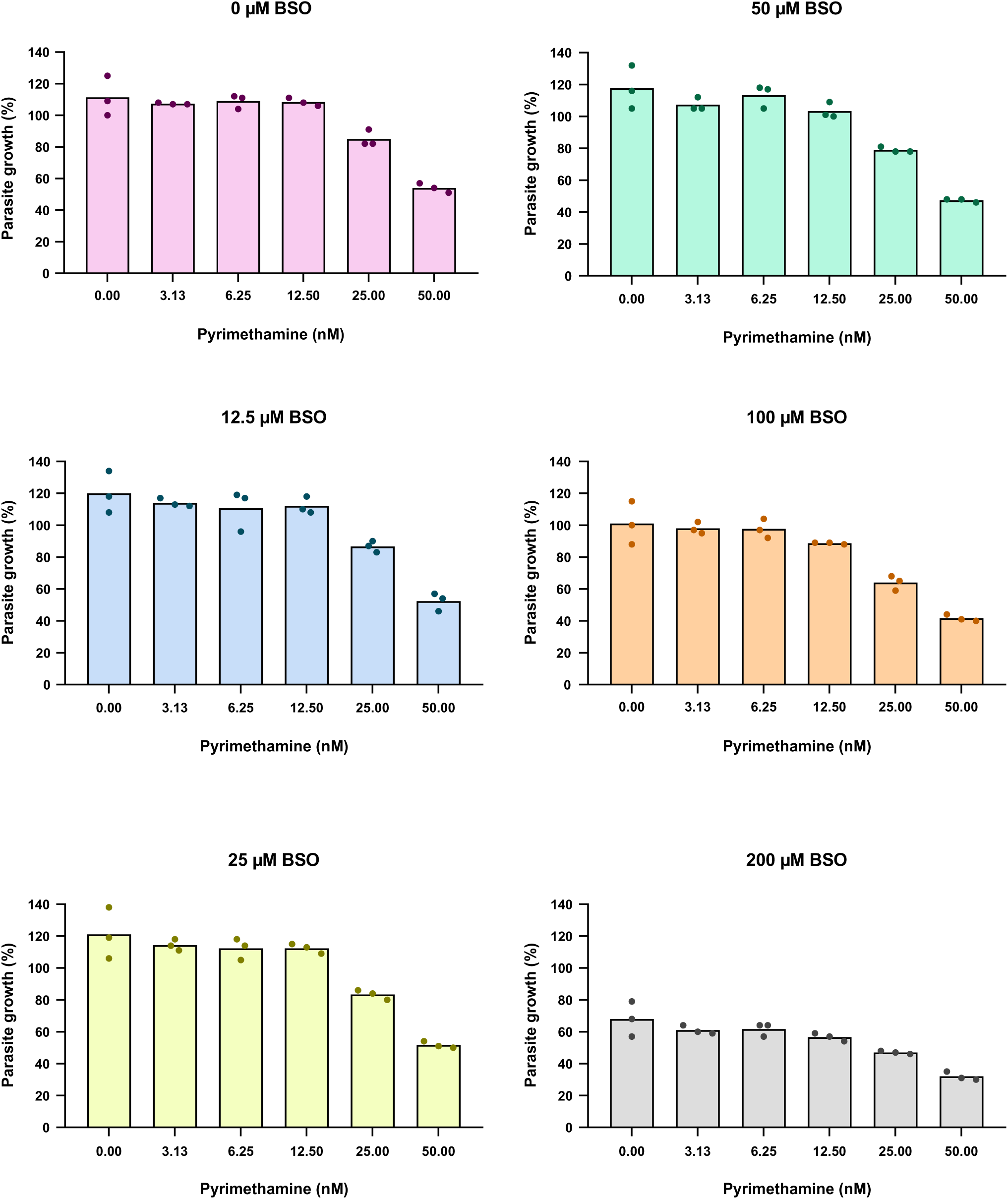
Treatment with Pyrimethamine and *L*-Buthionine-*S,R*-sulfoximine (BSO) as described in Suppl. Fig. 1. Corresponding to Figure 8C

**Supplemental Figure 20:**
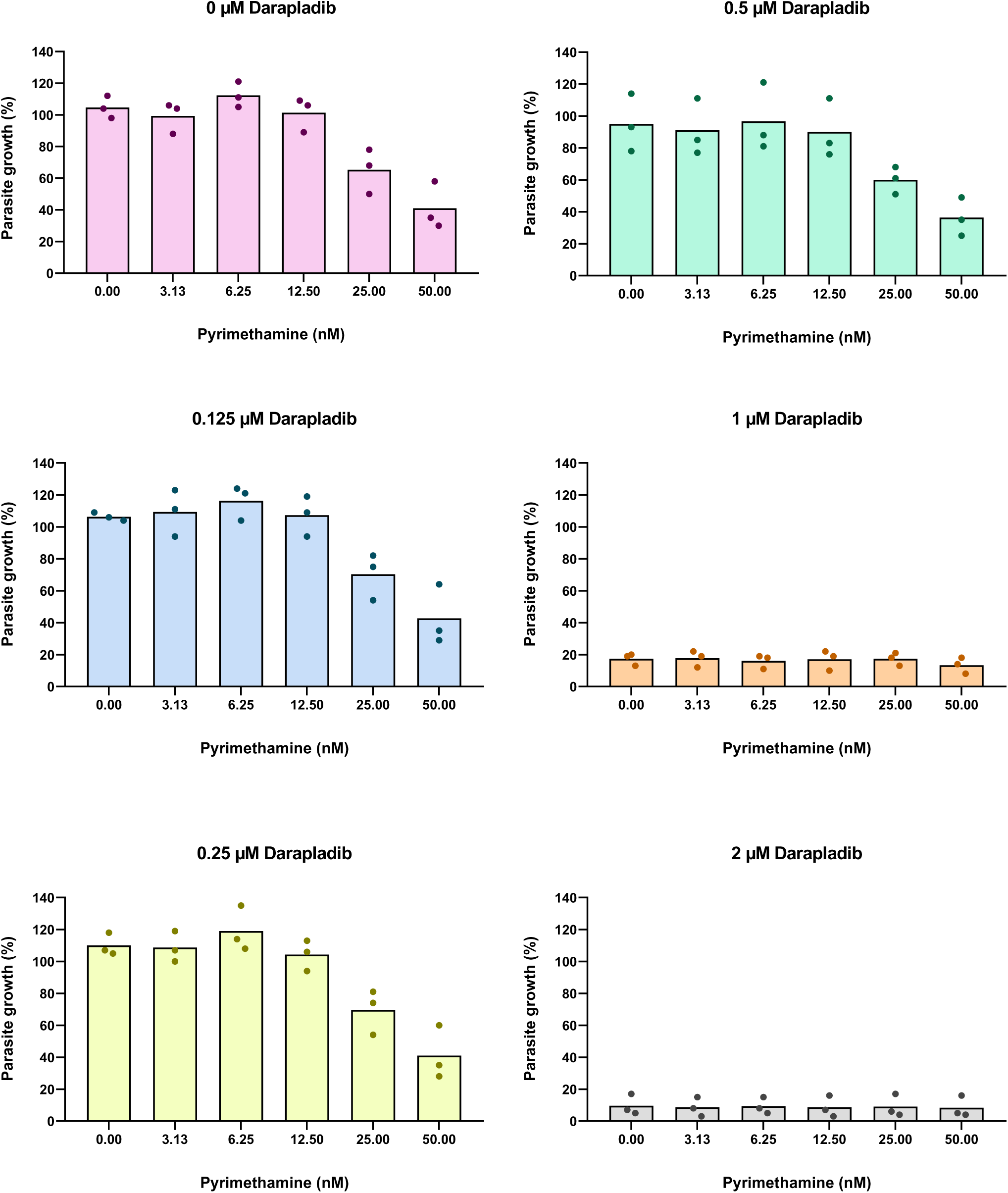
Treatment with Pyrimethamine and Darapladib as described in Suppl. Fig. 1. Corresponding to Figure 8D

**Supplemental Figure 21:**
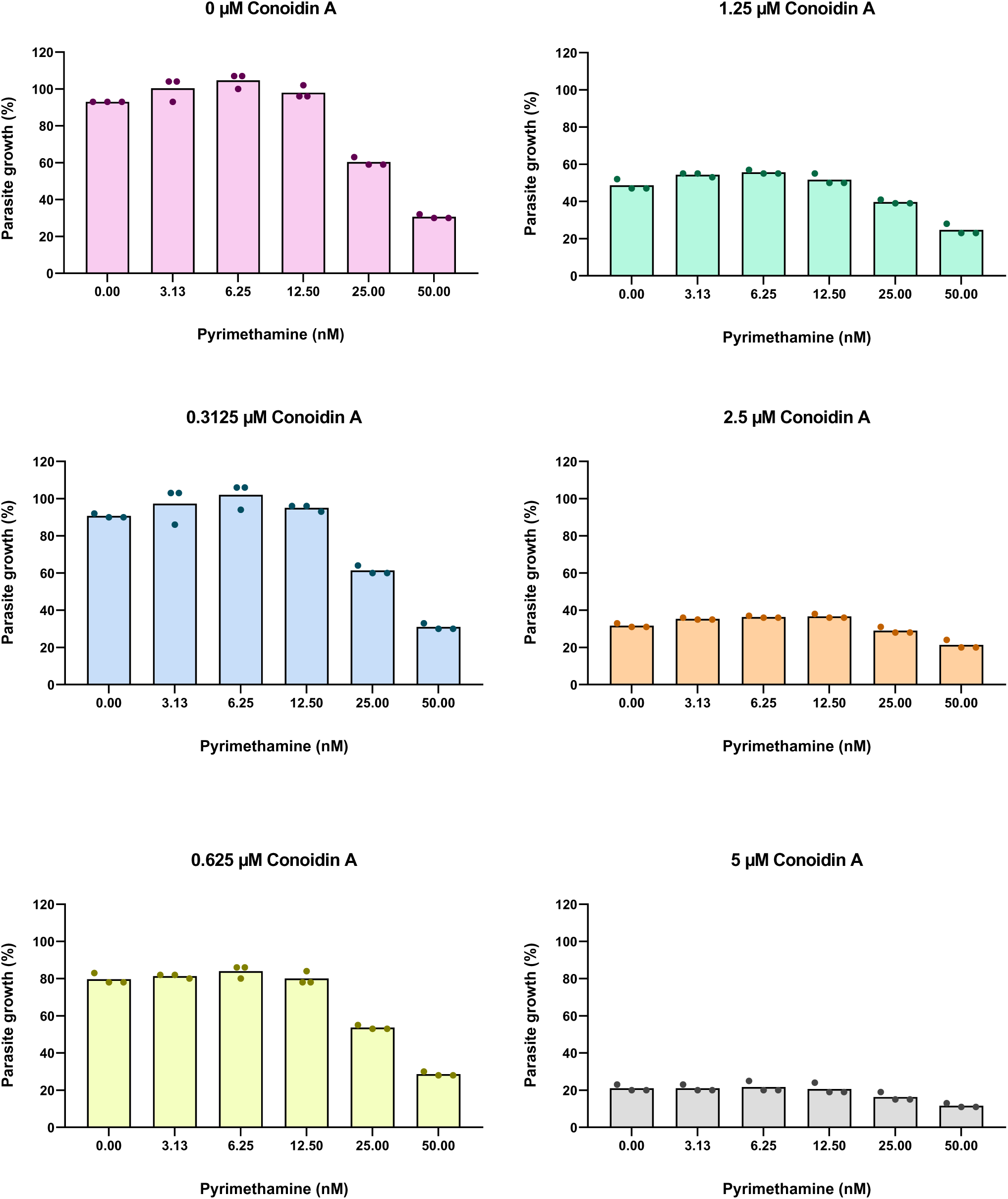
Treatment with Pyrimethamine and Conoidin A as described in Suppl. Fig. 1. Coresponding to Figure 8E

**Supplemental Figure 22:**
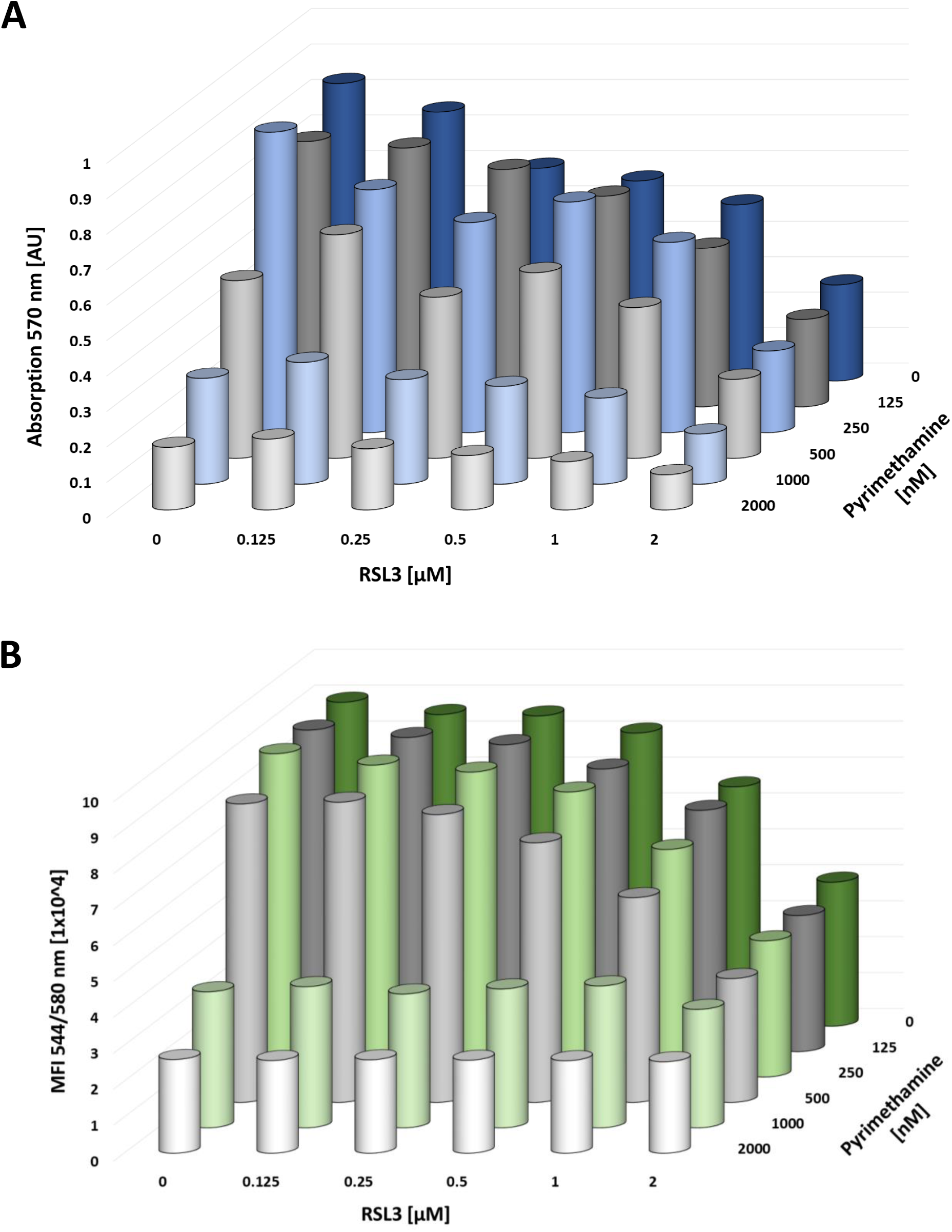
Lack of synergy between RSL3 and Pyrimethamine in the apicomplexan parasite *Toxoplasma gondii* **A.** Vero cells were infected with *T. gondii*, strain RH-LacZ. Cells were treated with RSL3 and Pyrimethamine at the indicated concentrations at 30 minutes post infection (p.i.). At 72 hours p.i., β-galactosidase activity was measured using chlorophenol red-β-D-galactopyranoside (CPRG) as a substrate. CPRG levels were measured by absorbance measurements at 570 nm, and represented as absorption units [AU]. Single-replicate values are displayed in Suppl. Fig. 23. **B.** Vero cells were infected with *T. gondii*, strain RH-pTUB-tdTomato RFP. Cells were treated with RSL3 and Pyrimethamine at the indicated concentrations at 30 minutes post infection (p.i.). Fluorescence as a measure of *T. gondii* RH-pTUB-tdTomato RFP growth was determined at 138 hours p.i. The mean fluorescence intensity (MFI) is presented as units of 1×10^4^. Single-replicate values are displayed in Suppl. Fig. 24.

**Supplemental Figure 23:**
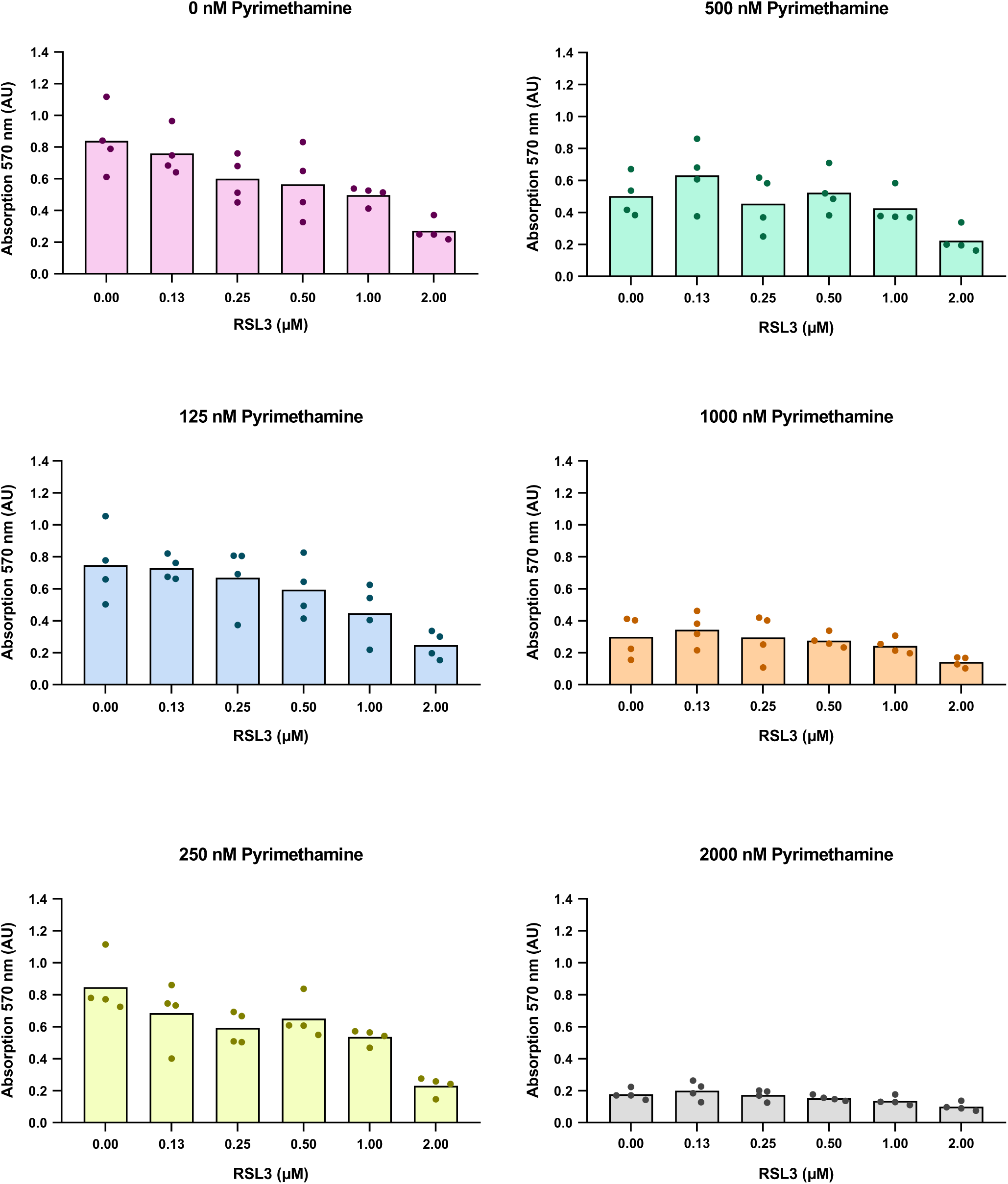
No synergy between RSL3 and Pyrimethamine in *T. gondii*, strain RH-LacZ. Values obtained from single biological replicates were plotted individually for each concentration of Pyrimethamine. Corresponds to **Suppl. Fig. 22A**

**Supplemental Figure 24:**
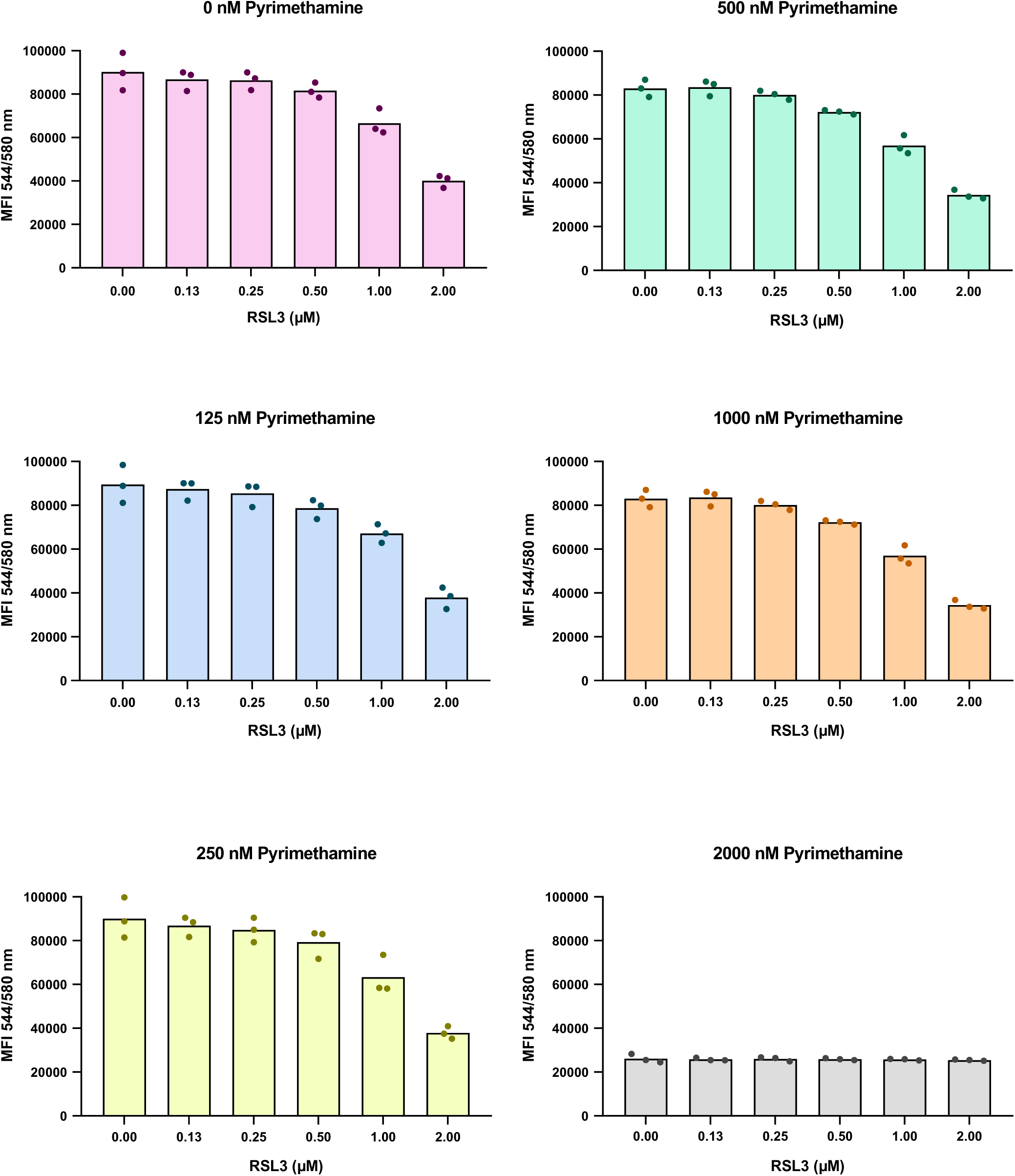
Lack of synergy between RSL3 and Pyrimethamine in *T. gondii*, strain RH-pTUB-tdTomato RFP. Values obtained from single biological replicates were plotted individually for each concentration of Pyrimethamine. Corresponds to **Suppl. Fig. 22B**

